# Lysine vitcylation is a novel vitamin C-derived protein modification that enhances STAT1-mediated immune response

**DOI:** 10.1101/2023.06.27.546774

**Authors:** Xiadi He, Yun Wei, Jiang Wu, Qiwei Wang, Johann S. Bergholz, Hao Gu, Junjie Zou, Sheng Lin, Weihua Wang, Shaozhen Xie, Tao Jiang, James Lee, John M. Asara, Ke Zhang, Lewis C. Cantley, Jean J. Zhao

## Abstract

Vitamin C (vitC) is a vital nutrient for health and also used as a therapeutic agent in diseases such as cancer. However, the mechanisms underlying vitC’s effects remain elusive. Here we report that vitC directly modifies lysine without enzymes to form vitcyl-lysine, termed “vitcylation”, in a dose-, pH-, and sequence-dependent manner across diverse proteins in cells. We further discover that vitC vitcylates K298 site of STAT1, which impairs its interaction with the phosphatase PTPN2, preventing STAT1 Y701 dephosphorylation and leading to increased STAT1-mediated IFN pathway activation in tumor cells. As a result, these cells have increased MHC/HLA class-I expression and activate immune cells in co-cultures. Tumors collected from vitC-treated tumor-bearing mice have enhanced vitcylation, STAT1 phosphorylation and antigen presentation. The identification of vitcylation as a novel PTM and the characterization of its effect in tumor cells opens a new avenue for understanding vitC in cellular processes, disease mechanisms, and therapeutics.

## INTRODUCTION

Humans, unlike most animals, have lost the ability to synthesize vitamin C (vitC) due to a mutation in the gene encoding the enzyme responsible for its production ^1^. As a result, humans rely on dietary intake to meet their vitC requirements. Low doses of vitC are essential for maintaining overall health and preventing diseases associated with vitC deficiency, such as scurvy ^2^. However, the use of high-dose vitC for treating diseases like cancer has been a topic of controversy over the last half-century ^3^.

Recent understanding of vitC pharmacokinetics through exploration of implications for both oral and intravenous administration has renewed interest and prompted further investigation into the clinical potential of vitC in cancer patients ^4–8^. Despite the growing interest in high-dose vitC utilization, the mechanisms behind its potential anti-cancer effects are not fully understood. One proposed mechanism is that vitC generates reactive oxygen species (ROS), which can selectively kill cancer cells. For example, several studies have shown that pharmacologic vitC can act as a prodrug for H_2_O_2_ formation, leading to the direct killing of cancer cells ^4–6, 9–13^. VitC has also been shown to selectively kill KRAS and BRAF mutant cancer cells via ROS accumulation in cells ^14–16^. Other reports have shown that vitC may exert its anti-tumor activity through DNA demethylation mediated by TET enzymes, where vitC functions as a cofactor ^17–21^.

VitC, also known as ascorbic acid (176 Da), exists in two main redox states: ascorbate anion (reduced form, 175 Da) and dehydroascorbic acid (DHA, oxidized form, 174 Da) ^22, 23^. Under physiological conditions, the predominant form of vitC is ascorbate anion (> 99%), which is more stable and biologically active than its oxidized counterpart, DHA (< 1%) ^22–24^. Previous studies have shown that, under low pH conditions (∼pH 2.0), DHA can undergo further oxidation to produce diketogulonate (DKG) ^25^. This oxidative process of DHA to DKG can lead to the formation of various chemical species that can modify specific amino acid residues, particularly cysteine or lysine residues, on proteins. These modifications, referred to as ascorbylations, occur primarily in plants, food products, and certain human tissues, such as the lens of the eyes ^26–29^. Protein post-translational modifications (PTMs) are increasingly appreciated for their crucial roles in physiological regulation due to their wide prevalence ^30–33^. Side-chain modifications involving the chemical alteration of specific amino acid residues, such as lysine, arginine, cysteine, serine, threonine, and tyrosine residues, within a protein are common forms of PTM ^32^. Some of the well-known side-chain modifications include acetylation, methylation, phosphorylation, and glycosylation, among others ^34–36^. Recent advancements in mass spectrometric technologies have led to the discovery of several novel side-chain modifications, such as cysteine carboxyethylation ^37^, glutamine dopaminylation ^38^, lysine lactylation ^39^, cysteine itaconate alkylation ^40^, and lysine aminoacylations ^41^. These modifications can modulate protein-protein interactions, enzymatic activity, protein stability, subcellular localization, and signaling pathways. By altering the chemical and physical properties of amino acids, side-chain modifications can have profound effects on protein structure and function.

In this study, we describe for the first time that ascorbate anion, the predominant form of vitC, can perform enzyme-free modifications on lysine residues in peptides and proteins under physiological conditions in a pH- and dose-dependent manner. We designate this novel PTM as “vitcylation” to distinguish it from the previously described ascorbylation induced by DHA. We provide evidence with cell-free biochemical and cellular biological assays to show that ascorbate anion is capable of directly modifying the ε-amine group of lysine in peptides and proteins. We further identify a broad range of vitcylated proteins by vitC, which cast lysine vitcylation as a form of side-chain modification that provides an enormous potential capacity for the cell to respond to fluctuation in vitC level. We demonstrate that vitC vitcylates the signal transducer and activator of transcription-1 (STAT1) at lysine-298 (K298) in cancer cells and that this modification results in increased STAT1 tyrosine 701 (Y701) phosphorylation and nuclear translocation. In seeking a mechanism underlying the relationship of STAT1 vitcylation and phosphorylation, we find that STAT1 K298 vitcylation impairs the interaction of STAT1 with protein tyrosine phosphatase non-receptor type 2 (PTPN2, also known as TC45), a STAT1 protein tyrosine phosphatase ^42, 43^, leading to increased STAT1 phosphorylation and upregulation of STAT1-mediated gene expression and activation of interferon (IFN) response pathway, thereby elucidating a pathway from STAT1 vitcylation by vitC to the immune responses both *in vitro* and in *vivo*. Our findings overall provide a molecular understanding of the role of vitC in protein modification and immune regulation with important implications for health and disease.

## RESULTS

### VitC directly modifies lysine to form vitcyl-lysine in cell-free systems in a pH- and dose-dependent manner

Previous studies have demonstrated that anhydride intermediates can react with lysine in proteins to form acyl-lysine protein modifications ^44^. For example, succinic anhydride reacts with lysine residues to form succinyl-lysine in proteins (succinylation) ^44, 45^ (**Figure S1A**). In addition, homocysteine (Hcy) can be converted to a reactive thioester intermediate, Hcy thiolactone (HTL), which in turn reacts with lysine residues to form homocysteinyl-lysine in proteins (N-homocysteinylation) ^46^ (**Figure S1B**). Anhydride intermediates and HTL possess a shared lactone structure that exhibits reactivity with lysine residues. We observed that vitC also contains a reactive lactone structure resembling those found in anhydride intermediates and HTL (**Figure 1A**). Based on this observation, we hypothesized that vitC may potentially modify the ε-amine group of lysine residues in peptides and proteins through its reactive lactone structure, resulting in the modified lysine, designated vitcyl-lysine (**Figure 1A**). To test this hypothesis, was generated three lysine-containing peptides as utilized in our previous study ^41^, and incubated them with vitC in a physiological pH condition (pH7.4). The Matrix-assisted laser desorption/ionization time-of-flight/time-of-flight mass spectrometry (MALDI-TOF/TOF MS) analysis unveiled a peptide mass shift of 175 Da (**Figure 1B**), indicating that it is the predominant form of vitC, ascorbate anion (175 Da), modified the ε-amine group of a lysine residue, forming vitcyl-lysine (**Figures 1A** and S1C).

**Figure 1.**
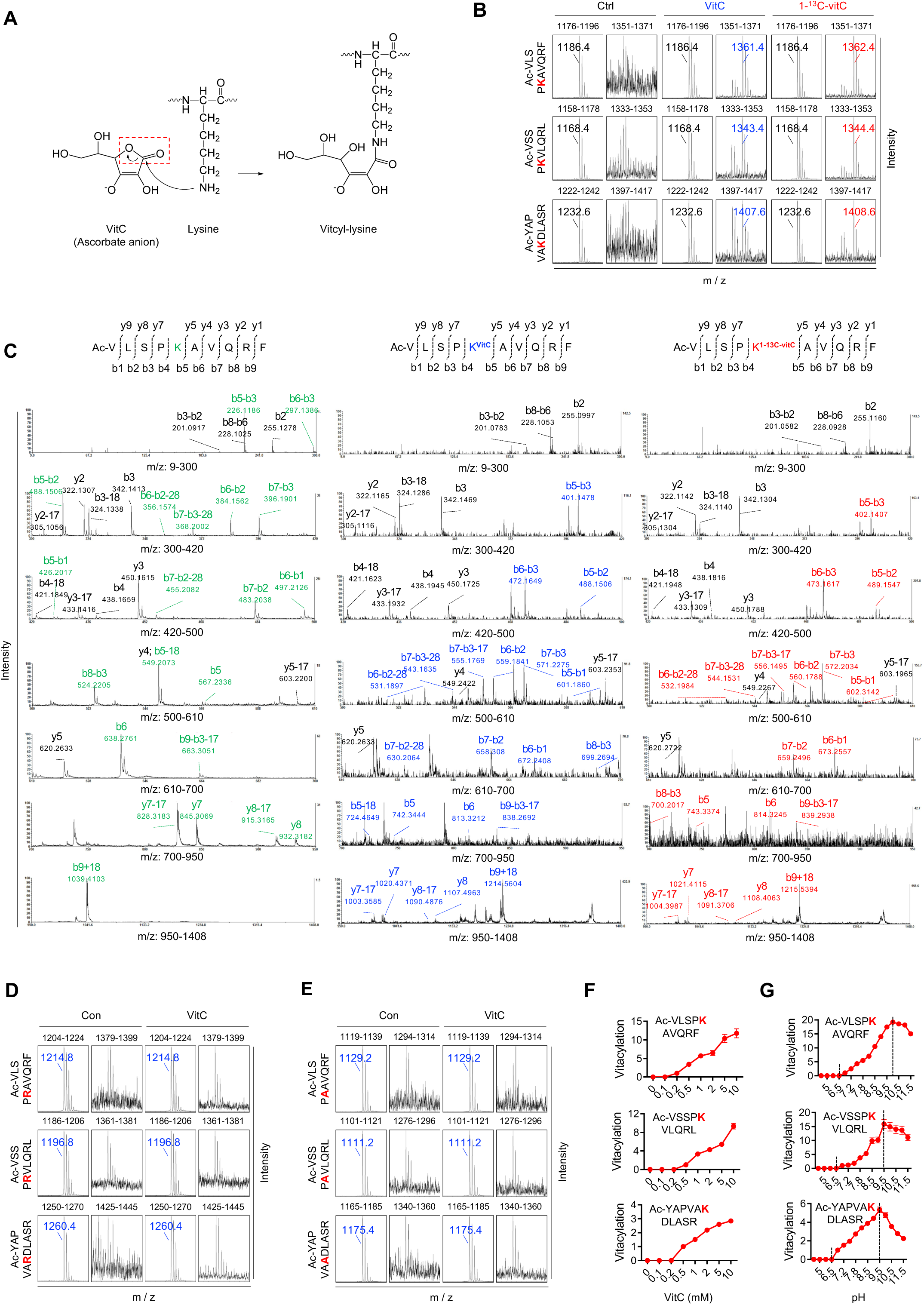
VitC modifies lysine residues of peptides to form vitcyl-lysine in cell-free systems. (A) Proposed mechanism of lysine vitcylation formation by ascorbate anion. The reactive lactone bond on ascorbate anion is circled. (B) Representative results of vitC-induced vitcylation formation *in vitro*. Synthetic lysine- containing peptides (sequences of the peptides were listed on the left of the spectrum, ‘Ac-’ means the N terminus of the peptide is protected by acetyl group, hereafter for MALDI-TOF/TOF MS detection, unless indicated otherwise) were incubated with either a vehicle, 2 mM vitC or 2 mM 1-^13^C-vitC at 37℃ for 3 hours. The formation of vitcylated peptides was detected by MALDI- TOF/TOF MS, with the m/z range of each spectrum displayed above the spectrum. The m/z of unmodified peptides and modified peptides were listed. (C) MS/MS spectrum of the unmodified peptide, vitcylated peptide, and 1-^13^C-vitcylated peptide (Ac-VLSPKAVQRF) detected by MALDI-TOF/TOF MS/MS. Lysine-containing unmodified fragments, vitcylated fragments, and 1-^13^C-vitcylated fragments are marked with green, blue, and red colors, respectively. (D and E) Synthetic arginine-containing peptides (D) and alanine-containing peptides (E) were incubated with either a vehicle or 2 mM vitC at 37℃ for 3 hours. The formation of vitcylated peptides was detected by MALDI-TOF/TOF MS. (F) Synthetic lysine-containing peptides were incubated with varying concentrations of vitC at 37℃ for 3 hours. The formation of vitcylated peptides was detected by MALDI-TOF/TOF MS. The relative vitcylation levels were quantified (right, n = 3). Data are represented as mean ± SEM. (G) Synthetic lysine-containing peptides were incubated with 2 mM vitC in different pH Tris-HCl buffer at 37℃ for 3 hours. The formation of vitcylated peptides was detected by MALDI- TOF/TOF MS, and the relative vitcylation levels were quantified (right, n = 3). Data are represented as mean ± SEM. See also Figure S1.

Previous studies have shown that under low pH conditions (∼pH 2.0), the oxidized form of vitC, DHA (174 Da), can undergo further oxidation to produce DKG, which in turn modifies cysteine or lysine residues of proteins in plants, food products, and the human lens with a mass shift ranging from 58 Da to 148 Da, and these modifications were termed “ascorbylations” (**Figure S1D**) ^26–29^. To determine whether DHA can form the modification under a physiological condition, we incubated the same three lysine-containing peptides with ascorbate anion or DHA at the same concentration (2 mM) and pH 7.4 in our cell-free system. Results showed that the incubation of the lysine-containing peptides with ascorbate anion, but not with DHA, formed vitcyl-lysine (**Figure S1E**). Thus we designate this ascorbate anion-induced lysing modification as “vitcylation” to distinguish it from the previously described ascorbylation induced by DHA.

To validate this ascorbate anion (vitC)-derived lysine modification, we incubated lysine-containing peptides with isotopic 1-^13^C-vitC (**Figure S1F**). Subsequent MALDI-TOF/TOF MS analyses showed that lysine 1-^13^C-vitcylation has a 176 Da mass shift (**Figure 1B**). Further MS/MS analysis of peptides with or without vitcylation or 1-^13^C-vitcylation confirmed that the lysine-containing vitcylated fragments have a 175 Da mass shift, and lysine-containing 1-^13^C-vitcylated fragments have a 176 Da mass shift (compared to the corresponding non-vitcylated fragments) (**Figures 1C, S1G, and S1H**).

To further confirm this new lysine-specific modification induced by vitC, we substituted the lysine residue in these peptides with arginine or alanine. Substituting lysine with arginine or alanine abolished the vitcylation induced by vitC (**Figures 1D and 1E**). Furthermore, we determined that this lysine modification by vitC displayed a sequence-specific preference, as the same modification was not observed in other lysine-containing peptides that we tested (**Figure S1I**). Together, these results demonstrate that vitC can directly modify peptides via vitcylation of lysine residues in our cell-free system, and this lysine vitcylation is at least partially sequence-specific.

We next sought to measure lysine vitcylation with a range of vitC concentrations from 0.1 mM to 10 mM in our cell-free system (plasma vitC concentrations greater than 10 mM are easily achieved in humans without significant toxicity) ^47, 48^, as well as with a pH scale from pH 4.0 to pH 11.5 to encompass the physiological pH across different subcellular compartments from 6.5 to 8.2 ^49–51^. The vitC dose titration (0.1-10 mM) showed that increasing concentrations of vitC resulted in increasing levels of vitcylation with an EC_50_ value of approximately 2 mM (**Figure 1F**). We then performed a pH titration of vitcylation on lysine-containing peptides with vitC at 2 mM. Notably, we observed a steep elevation of vitcylation levels from pH 7.0 to pH 9.5-10.0 followed by a quick decline at higher pH (**Figure 1G**). Together, our data indicate that lysine vitcylation is a hitherto unknown modification by vitC in the ascorbate anion form in our cell-free and enzyme-free systems in a dose-, pH-, and sequence-dependent manner.

### VitC modifies lysine in cellular proteins to form lysine-vitcylated proteins

We subsequently conducted vitcylation with the whole cell protein mix isolated and purified from E0771 cell lysates by acetone precipitation ^52^. Reaction mixtures of vitC (1 mM) with purified proteins were established in two different pH conditions, pH 7.2 and pH 8.0, respectively. High-performance liquid chromatography (HPLC)–MS/MS analysis revealed that vitC generated more abundant lysine vitcylation in proteins at pH 8.0 than at pH 7.2 (**Figure S2A; Tables S1-S3**).

We further show that vitC treatment leads to a substantial rise in the cellular concentration of vitC within both cultured human cancer cells (Cal-51) and murine cancer cells (E0771 and PP) in a dose-dependent manner (**Figures 2A**). PP is a recently developed syngeneic mouse breast tumor model driven by the concurrent loss of PTEN and p53 in the Zhao lab ^53^. To investigate whether vitC can modify cellular proteins within intact cells, we treated human Cal-51 and mouse E0771 cells with 2 mM vitC and prepared cell lysates for analysis of vitcylation of cellular proteins. Mass spectrometry analysis identified 573 and 1450 proteins with vitcylations in Cal-51 and E0771 cells, respectively, with 94 shared between both cell lines (**Figures 2B and 2C; Tables S4-S6**). Further bioinformatic analysis revealed that the vitcylated proteins are widely distributed across different subcellular locations with potential biological roles in multiple cellular processes and signaling pathways (**Figures 2D-2G and S2B-S2E**). Consistent with the sequence-specific feature of vitC-mediated vitcylation in lysine-containing synthetic peptides (**Figure S1I**), we found enriched lysine vitcylation motifs in proteins, although we did not identify a strong consensus sequence for vitcylation (**Figure S2F)**.

**Figure 2.**
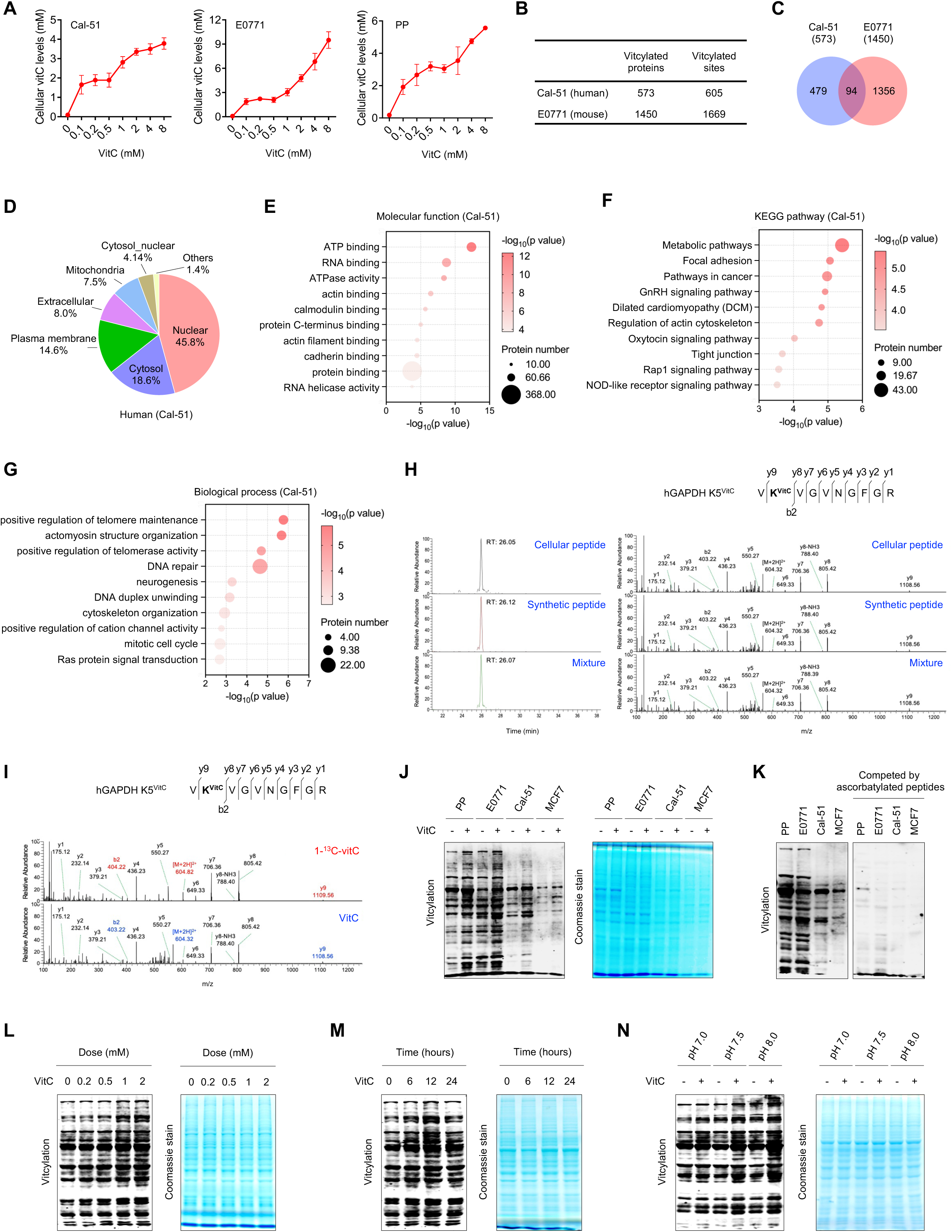
VitC induces lysine vitcylation on cellular proteins. (A) Intracellular vitC levels were measured in Cal-51, E0771, and PP cells cultured in different concentrations of vitC for 12 hours (n = 3). Data are represented as mean ± SEM. (B) Numbers of vitcylated proteins and sites identified in Cal-51 (human) and E0771 cells (mouse) are summarized. (C) Venn diagram of shared vitcylation proteins identified in Cal-51 cells (human) and E0771 cells (mouse). (D) Subcellular locations of lysine vitcylated proteins identified in Cal-51 cells. The locations are classified into nuclear, cytosol, plasma membrane, extracellular, mitochondrial, cytosol_nuclear, and other compartments. (E) Top ten gene ontology molecular function enrichment of vitcylated proteins identified in Cal- 51 cells. (F) Top ten KEGG-based enrichment of lysine vitcylated proteins identified in Cal-51 cells. (G) Top ten gene ontology biological process enrichment of vitcylated proteins identified in Cal- 51 cells. (H) Extracted ion chromatograms (left) and MS/MS spectra (right) from HPLC-MS/MS analysis of a vitcylated peptide (human GAPDH, K5) derived from Cal-51 cells (cellular peptide), its *in vitro* generated counterpart (synthetic peptide), and their mixture. The b ion refers to the N-terminal parts of the peptide, and the y ion refers to the C-terminal parts of the peptide (hereafter for HPLC- MS/MS analysis). (I) Extracted MS/MS spectra from HPLC-MS/MS analysis of 1-^13^C-vitcylated peptides and vitcylated peptides (human GAPDH, K5) derived from Cal-51 cells (in cells, the lysine-containing 1-^13^C-vitcylated fragments and vitcylated fragments were marked by red and blue colors, respectively). (J) Intracellular lysine vitcylation levels were measured from PP, E0771, Cal-51, and MCF7 cells cultured in medium containing a vehicle or vitC for 12 hours (2 mM vitC for PP and E0771 culture, 0.5 mM vitC for Cal-51 and MCF7 culture). Protein levels in each sample were normalized by coomassie staining, hereafter for global vitcylation detection. (K) Vitcylation signals of indicated cells were competed off by vitcylated peptide (Ac- VLSPKAVQRF peptide pre-incubated with 2 mM vitC at 37℃ for 3 hours). (L) Intracellular lysine vitcylation levels were measured from E0771 cells cultured in medium containing different concentrations of vitC for 12 hours. (M) Intracellular lysine vitcylation levels were measured in E0771 cells cultured in medium containing 2 mM vitC for the indicated times. (N) Intracellular lysine vitcylation levels were measured from E0771 cultured in medium containing a vehicle or 2 mM vitC for 12 hours under different pH conditions. See also Figure S2 and Tables S1-S6.

We next determined whether the lysine-vitcylation induced by vitC in cellular proteins is the same modification observed in synthetic peptides in our cell-free system described above (**Figure 1**). To achieve this, we employed HPLC–MS/MS to separate and analyze the vitcylated peptides from E0771 cells treated with vitC and compared them with lysine-containing synthetic peptides treated with vitC in the cell-free system. Indeed, each pair of peptides co-eluted from HPLC had comparable MS/MS spectra (**Figures 2H, S2G, and S2H**). Treatment of the cells with isotopic 1- ^13^C-vitC followed by MS/MS analysis further validates that vitC induces lysine vitcylation in proteins within cells (**Figures 2I, S2I, and S2J**). Overall, these results demonstrate that vitC treatment leads to lysine vitcylation in cellular proteins.

To further validate lysine vitcylation in cells, we developed polyclonal antibody against vitcyl-lysine and confirmed its specificity by dot-blot assay (**Figure S2K**). Western blot (WB) analysis using this anti-vitcylation antibody detected specific bands in cell lysates prepared from vitC-treated cancer cells (**Figure 2J**). Notably, these bands could be outcompeted by vitcylated peptides in a WB experiment (**Figure 2K**), suggesting that this antibody indeed recognizes lysine vitcylation and supports the occurrence of vitcylation in cells. Further WB analysis with the anti-vitcylation antibody demonstrated that lysine vitcylation levels in cells respond to vitC in a dose-, time-, and pH-dependent manner (**Figures 2L-2N**). Taken together, these results demonstrate that vitC in the form of ascorbate anion can modify lysine in proteins in human and murine cells with vitcylatiion.

### STAT1 K298 vitcylation enhances STAT1 phosphorylation and activation

We proceeded to explore a potential functional role of lysine vitcylation in cells. Expression analysis across a panel of 4,604 cancer- and immune-related genes from E0771 cells treated with vitC revealed that the expression of genes within gene ontology (GO) terms relating to immune activities and inflammatory responses were upregulated by vitC (**Figure 3A**). We further conducted gene set enrichment analysis (GSEA) and found that the top-ranked genes and their associated cellular processes in vitC-treated cells were related to the ‘IFNγ response’, ‘IFNα response’ and ‘inflammatory response’ (**Figures 3B, S3A, and S3B**).

**Figure 3.**
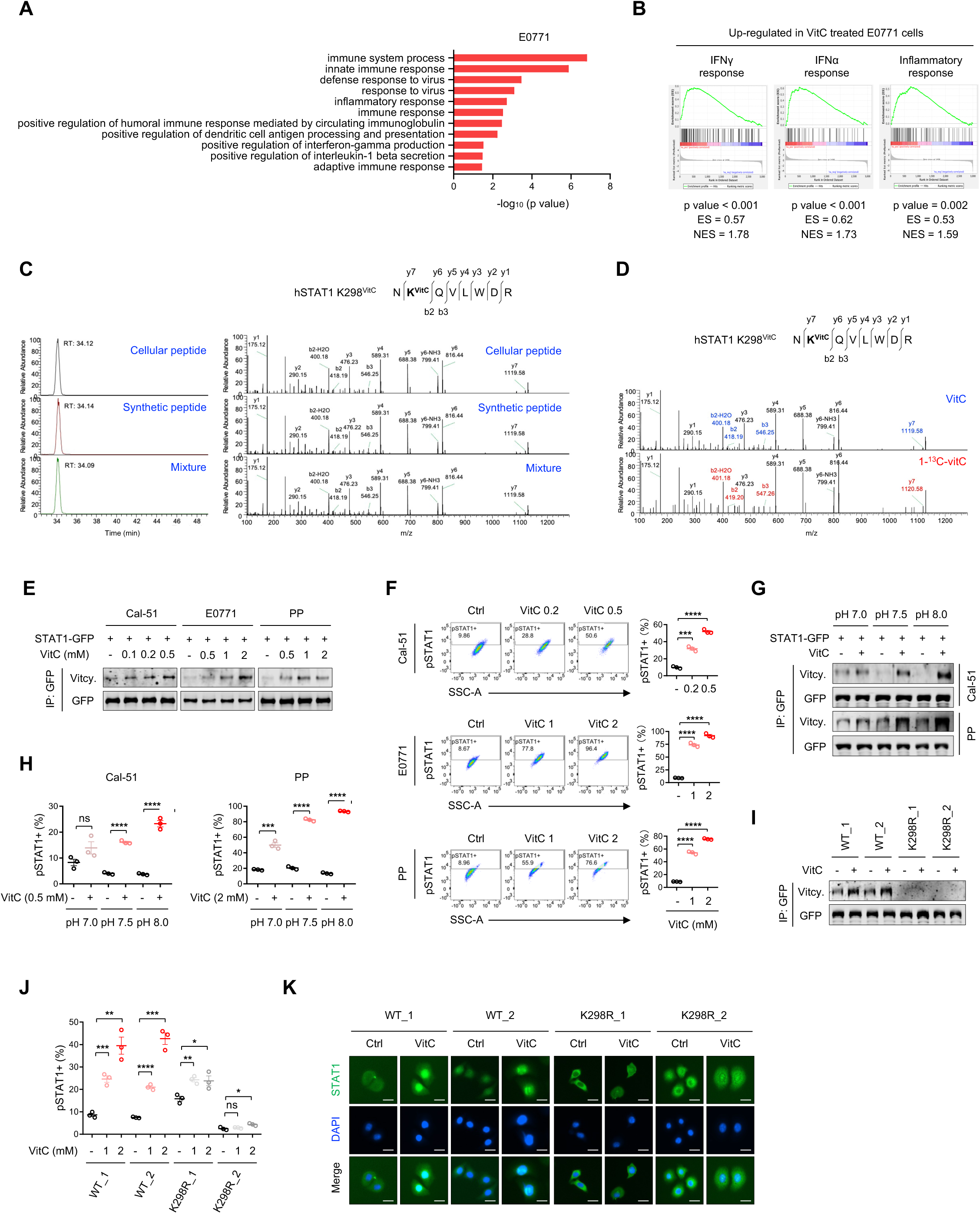
Vitcylation of STAT1 K298 regulates the phosphorylation and activation of STAT1. (A and B) Top-ranked upregulated GO terms (A) and upregulated GSEA signatures (B) in E0771 cells treated with 1 mM vitC for 2 days (n = 2). (C) Extracted ion chromatograms (left) and MS/MS spectra (right) from HPLC-MS/MS analysis of a vitcylated peptide (human STAT1, K298) derived from Cal-51 cells (cellular peptide), its *in vitro* generated counterpart (synthetic peptide) and their mixture. (D) Extracted MS/MS spectra from HPLC-MS/MS analysis of vitcylated peptide (upper) and 1-^13^C-vitcylated peptide (lower) (human STAT1, K298) derived from Cal-51 cells. Lysine-containing vitcylated fragments and 1-^13^C-vitcylated fragments are marked by blue and red colors, respectively. (E) STAT1 vitcylation levels were measured from STAT1-GFP expressing cells (Cal-51 and PP cells) cultured in different pH mediums with or without vitC for 12 hours (2 mM vitC for PP cell culture, 0.5 mM vitC for Cal-51 cell culture). (F) pSTAT1 flow cytometric analysis of Cal-51 and PP cells cultured in different pH mediums with or without vitC for 2 days (2 mM vitC for PP cell culture, 0.5 mM vitC for Cal-51 cell culture, n = 3). Data are represented as mean ± SEM. ***p < 0.001, ****p < 0.0001. (G) STAT1 vitcylation levels were measured from STAT1-GFP expressing cells (Cal-51 and PP cells) cultured in different pH mediums with or without vitC for 12 hours (2 mM vitC for PP cell culture, 0.5 mM vitC for Cal-51 cell culture). (H) pSTAT1 flow cytometric analysis of Cal-51 and PP cells cultured in different pH mediums with or without vitC for 2 days (2 mM vitC for PP cell culture, 0.5 mM vitC for Cal-51 cell culture, n = 3). Data are represented as mean ± SEM. ***p < 0.001, ****p < 0.0001. (I) Measurement of STAT1-WT and STAT1-K298R vitcylation levels in PP-sgSTAT1_1 cell re- expressing STAT1-WT-GFP or STAT1-K298R-GFP cultured in 2 mM vitC-containing or control medium for 12 hours. (J) pSTAT1 flow cytometric analysis of PP-sgSTAT1_1 cell re-expressing STAT1-WT-GFP or STAT1-K298R-GFP cultured in different concentrations of vitC for 2 days (n = 3). Data are represented as mean ± SEM. *p < 0.05, **p < 0.01, ***p < 0.001, ****p < 0.0001. (K) Nuclear translocation of STAT1 in PP-sgSTAT1_1 cell re-expressing STAT1-WT-GFP or re- expressing STAT1-K298R-GFP treated with 2 mM vitC for 2 days was assessed by immunofluorescence (scale bar, 50 μM). See also Figure S3 and Table S4.

Since STAT1 activation initiates most IFN response transcription programs ^54^, and our initial analysis of vitC-induced vitcylated proteins revealed that STAT1 lysine-298 (K298) in cells was vitcylated upon vitC treatment (**Table S4; Figure S3C**), we decided to further investigate STAT1 modification in response to vitC treatment. STAT1 K298 is a crucial regulatory site for STAT1 activity and is an evolutionarily conserved site in vertebrate animals (**Figure S3D**) ^55–57^. We performed more detailed analyses specifically on the vitcylation of STAT1 K298 in human Cal-51 and murine E0771 cell lines. HPLC-MS/MS analysis was used to compare the vitcylated peptides derived from STAT1 K298 in vitC-treated cells with those derived from vitC-treated synthetic peptides containing STAT1 K298. Notably, each pair of peptides co-eluted in HPLC and had comparable MS/MS spectra (**Figures 3C and S3E**). Treatment of the cells with isotopic 1-^13^C- vitC followed by MS/MS analysis further confirmed that vitC modified STAT1 K298 to form vitcyl-K298 in both human and mouse cells (**Figures 3D and S3F**).

We next investigated whether STAT1 vitcylation contributed to the increased cellular immunity and inflammatory response seen upon vitC treatment. Since phosphorylation of STAT1 at tyrosine 701 (pSTAT1) is crucial for its nuclear translocation and subsequent IFN responses in cells ^58, 59^, we hypothesized that vitcylation of STAT1 may impact its phosphorylation. To test this, we examined the correlation between STAT1 vitcylation and phosphorylation in cancer cells treated with vitC. STAT1 vitcylation was assessed by pull-down of GFP-tagged STAT1 (STAT1-GFP) expressed in cells with GFP-antibody followed by WB analysis with anti-vitcylation antibody. We observed that vitC treatment dose-dependently increased both vitcylation and phosphorylation levels of STAT1 in Cal-51, E0771, and PP cells (**Figures 3E and 3F**). Moreover, the levels of both vitcylation and phosphorylation in STAT1 were found to increase in response to vitC treatment, exhibiting a pH-dependent pattern ranging from pH 7.0 to 8.0 (**Figures 3G and 3H**). This observation is consistent with our previous finding that vitC induces pH-dependent vitcylation in our cell-free system (**Figure 1G**). Additionally, we confirmed enhanced STAT1 nuclear translocation in cells upon vitC treatment (**Figure S3G).** These results suggest that vitC-induced lysine vitcylation of STAT1 is closely associated with its phosphorylation and subsequent nuclear translocation, providing a potential mechanism for the observed enhanced cellular immunity and inflammatory response.

To further study the role of STAT1 vitcylation in the regulation of STAT1 phosphorylation and activation, we generated STAT1-null PP tumor cells via CRISPR/Cas9-mediated gene editing (**Figure S3H**), and reintroduced either wild-type STAT1 (STAT1-WT) or a vitcylation defective K298R mutant STAT1 (STAT1-K298R) (**Figure S3I**). Notably, adding back STAT1-WT, but not STAT1-K298R, into STAT1-null PP cells restored STAT1 vitcylation, as well as enhanced STAT1 phosphorylation and STAT1 nuclear accumulation upon vitC treatment (**Figures 3I-3K**). These results further validate that STAT1 vitcylation modulates phosphorylation and activation of STAT1 in tumor cells.

### STAT1 K298 vitcylation prevent STAT1 from dephosphorylation by its phosphatase PTPN2

We next sought to further understand the molecular mechanism underlying the relationship between STAT1 phosphorylation and vitcylation. A set of STAT1 gain-of-function (GOF) mutations has been identified as the genetic etiology of chronic mucocutaneous candidiasis (CMC), an autoimmune disorder ^60^. This GOF mechanism involves impaired STAT1 dephosphorylation that results in STAT1 hyperphosphorylation at Y701 in response to type I and II IFNs stimulation ^60, 61^. Interestingly, one of these GOF mutations is K298N, and cells with STAT1-K298N exhibit higher pSTAT1 levels both in basal conditions and after IFNγ stimulation ^55^. Furthermore, many STAT1 GOF mutations are located close to K298 in the STAT1 secondary/tertiary structure (**Figure 4A**). Therefore, we hypothesized that STAT1-K298 vitcylation may prevent STAT1 from dephosphorylation by its phosphatase. To test this idea, we performed co-immunoprecipitation of STAT1 with its phosphatase PTPN2, a STAT1 protein tyrosine phosphatase ^42, 43^, in the presence or absence of vitC treatment. Indeed, although STAT1 was able to co-immunoprecipitate PTPN2, vitC treatment abolished this association (**Figure 4B)**. In addition, although vitC treatment has little effect on the association of STAT1 with its protein kinase JAK1 (**Figure S4A**) ^54^, STAT1 phosphorylation in vitC-treated cells was sustained following the treatment with tyrosine kinase inhibitor staurosporine upon IFNγ stimulation (**Figures 4C and S4B**). Collectively, these results demonstrate that STAT1 vitcylation enhances STAT1 phosphorylation by impairing STAT1 dephosphorylation by its phosphatase.

**Figure 4.**
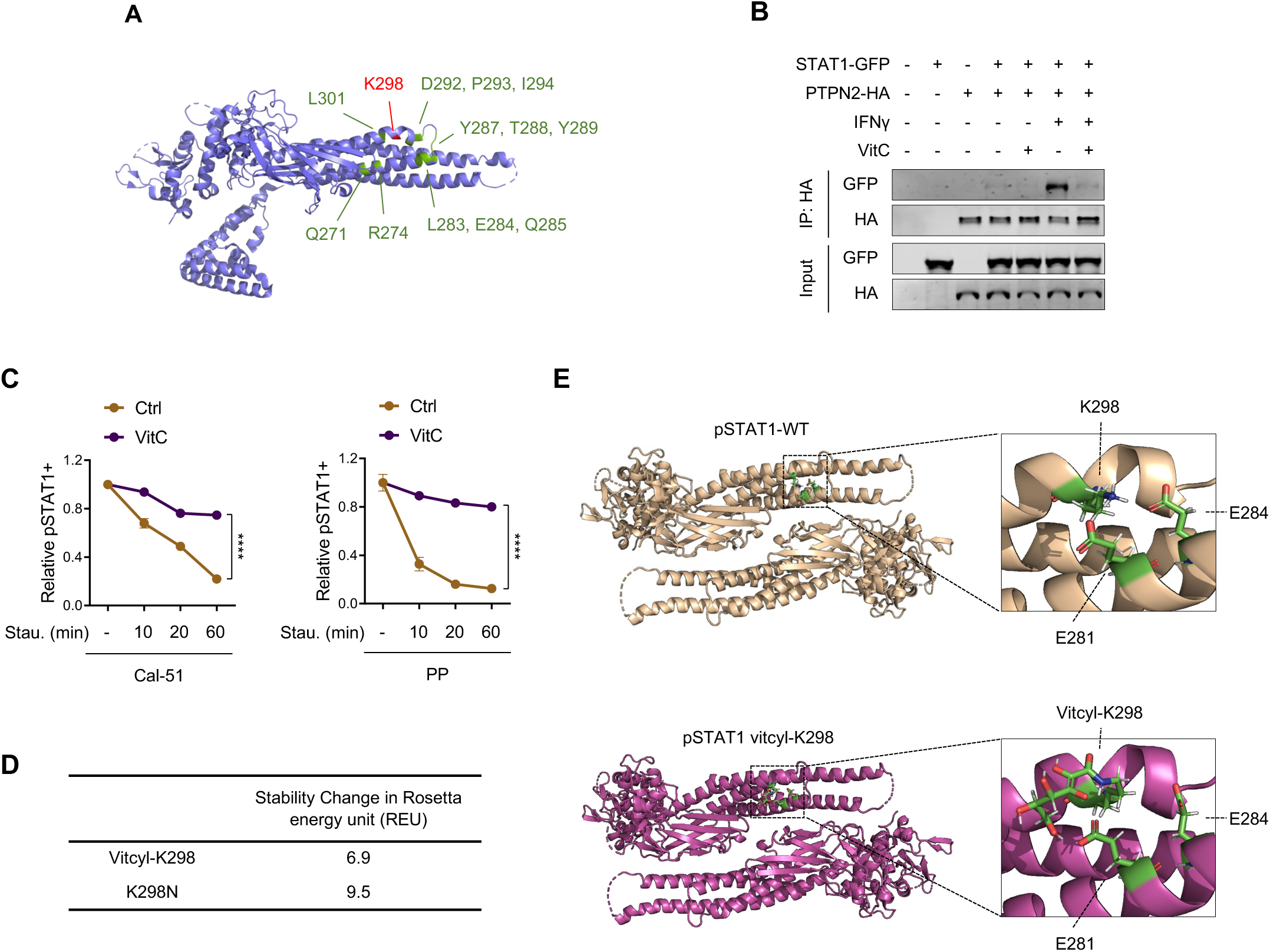
Vitcylation of STAT1 K298 prevents its dephosphorylation by PTPN2. (A) Ribbon representation of human STAT1. The K298 site and several gain-of-function mutation sites were marked by red and blue colors, respectively. The side chain of K298 was shown. (B) HeLa cells co-expressing STAT-GFP and PTPN2-HA were treated with vehicle or 300 μM vitC for 1 day, followed by stimulation with 100 ng/ml IFNγ for 15 min. Interaction between STAT1 and PTPN2 was assayed by co-immunoprecipitation. (C) Cells were pretreated with vehicle or vitC (0.2 mM vitC for Cal-51 cell culture, 1 mM vitC for PP cell culture) for 2 days, then cells were stimulated with 100 ng/ml IFNγ for 15 min followed by incubation with 1 μM staurosporine for indicated times. The relative pSTAT1+ populations were measured by flow cytometry immediately (n = 3). Data are represented as mean ± SEM. ****p < 0.0001. (D) Stability change in Rosetta energy unit (REU) of STAT1 caused by K298 vitcylation and K298N mutation as determined by the Rosetta atom energy function model system. (E) Structures of wild-type and K298 vitcylated pSTAT1 in the antiparallel dimer conformation from the last snapshot of MD simulation. Vitcyl-K298 loses the salt bridges of K298/E281 and K298/E284 in STAT1. See also Figure S4.

Previous studies have shown that IFNγ stimulation results in the formation of parallel pSTAT1 homodimers and their recruitment to gamma interferon activation sites (GAS DNA elements), both key events in the IFN signaling pathway ^62–64^. Interestingly, recent studies have shown that a rapid conformational rearrangement of pSTAT1 dimers from a parallel to an antiparallel dimer conformation seems to be required for binding to PTPN2 ^65–67^. To assess whether STAT1-K298 modifications, such as the K298-vitcylation and K298N mutation, impede the transition to the antiparallel dimer conformation, we employed the Rosetta atom energy function system for biomolecular modeling to identify and analyze the structural conformation changes with pSTAT1- K298 modifications. Stability changes in Rosetta energy unit (REU) showed that both K298 vitcylation and K298N had significant destabilizing effects on the antiparallel dimer form of STAT1 (with REU 6.9 and 9.5 respectively) (**Figure 4D**). The total stability changes were further decomposed into different energy terms in the Rosetta scoring function (**Figure S4C**). Both vitcyl- K298 and K298N mutation have lost the salt bonds that of K298 forms with E281 and E284 in their antiparallel dimer conformation (**Figures 4E and S4D**). Together, these data suggest that both STAT1-vitcyl-K298 and STAT1-K298N have impaired rearrangement of pSTAT1 parallel dimers (required for DNA binding) to an antiparallel dimer conformation (required for dephosphorylation by PTPN2), resulting in increased STAT1 phosphorylation and activation. Interestingly, while STAT1-K298N is a GOF genetic mutation and STAT1-K298-vitcylation is a chemical modification by vitC, they share a common underlying molecular mechanism for their enhancement of the immune response.

### STAT1 K298 vitcylation enhances MHC/HLA class I expression and immunogenicity in tumor cells

Since STAT1 phosphorylation and activation lead to IFN-mediated antigen processing and presentation in cells ^68^, we performed real-time quantitative polymerase chain reaction (RT-PCR) expression analysis of genes associated with antigen processing and presentation, and flow cytometry analyses of major histocompatibility complex (MHC/HLA) class I expression. We found that vitC treatment significantly upregulated the expression of multiple antigen processing and presentation genes, such as *Tap1, Lmp2, H2k1, B2m* and *Irf1* in PP cells and *HLA-B, TAP1, TAP2, LMP2* and *B2M* in Cal51 cells (**Figure S5A)**. Moreover, we observed a dose- and pH-dependent increase in surface protein levels of MHC class I in PP cells and HLA class I in Cal-51 cells following vitC treatment (**Figures 5A and 5B**). We further show that vitC-induced MHC/HLA class I expression was abolished by the loss of STAT1, and overexpression of STAT1- WT, but not the STAT1-K298R mutant, rescued vitC-induced MHC/HLA class I expression in STAT1-null PP cells (**Figures 5C and S5B**). These results suggest that STAT1 vitcylation induced by vitC contributes to the activation of STAT1 and the upregulation of MHC/HLA class I components in these cells.

**Figure 5.**
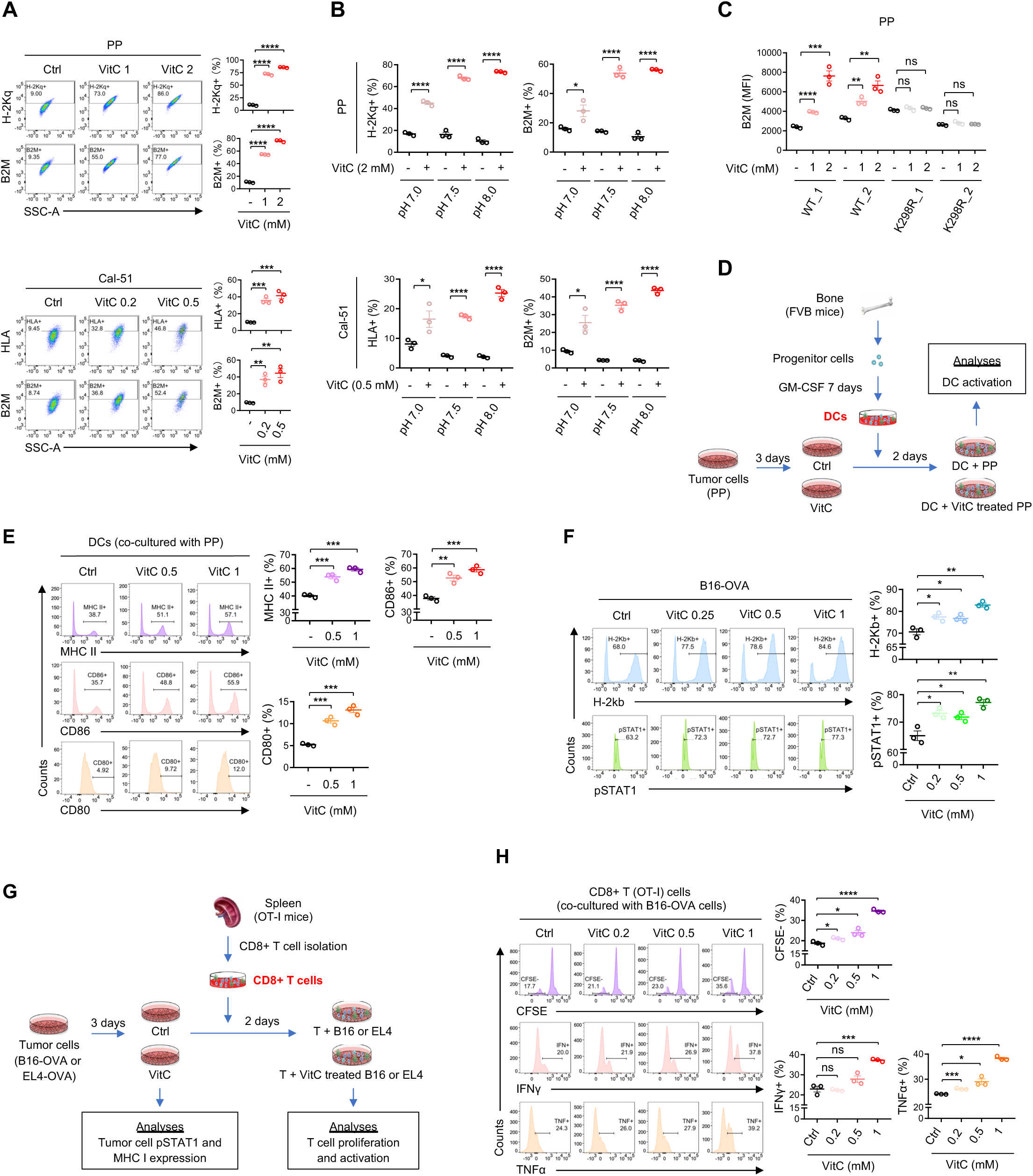
Vitcylation of STAT1 K298 enhances the expression of MHC/HLA class I and promotes immunogenicity in tumor cells. (A) Representative flow cytometry plots (left) and quantifications (right) of MHC class I expression on PP (upper) and Cal-51 cells (lower) cultured with different concentrations of vitC for 2 days (n=3). Data are represented as mean ± SEM. **p < 0.01, ***p < 0.001, ****p < 0.0001. (B) Flow cytometric analysis of MHC class I expression on PP (upper) and Cal-51 cells (lower) cultured in different pH mediums with or without vitC for 2 days (2 mM vitC for PP cell culture, 0.5 mM vitC for Cal-51 cell culture) (n = 3). Data are represented as mean ± SEM. *p < 0.05, ****p < 0.0001. (C) Flow cytometric analysis of MHC class I expression on PP-sgSTAT1_1 cell overexpressed with STAT1-WT-GFP (2 single clones) or STAT1-K298R-GFP (2 single clones) cultured with different concentrations of vitC for 2 days (n = 3). Data are represented as mean ± SEM. **p < 0.01, ***p < 0.001, ****p < 0.0001. (D) Workflow for co-culturing of vitC-treated PP cells with bone marrow-derived DCs. (E) Flow cytometry analysis of DCs co-cultured with vitC-pretreated PP cells. DCs (CD45^+^ CD11c^+^) were plotted and quantifications as MHC II^+^, CD86^+^ and CD80^+^ to identify DCs activity (n = 3). Data are represented as mean ± SEM. **p < 0.01, ***p < 0.001. (F) Flow cytometric analysis of H-2Kb and pSTAT1 expression on B16-OVA cells treated with different doses of vitC for 3 days (n = 3). Data are represented as mean ± SEM. *p < 0.05, **p < 0.01. (G) Workflow for co-culturing of vitC-treated B16-OVA or EL4-OVA cells with OT-I mice spleen- derived CD8^+^ T cells. (H) Flow cytometric analysis of CD8^+^ T (OT-I) cells co-cultured with B16-OVA cells pretreated with 2 mM vitC. T cells (CD45^+^ CD3^+^ CD8^+^) proliferation and activity were quantified as CFSE^-^ and IFNγ^+^, TNFα^+^ cells, respectively (n = 3). Data are represented as mean ± SEM. *p < 0.05, ***p < 0.001, ****p < 0.0001. See also Figure S5.

Previous reports have shown that many of the physiological functions of vitC involve its ability to increase the levels of ROS and act as a cofactor for ten-eleven translocation (TETs) enzymes and hypoxia-inducible factor 1-alpha (HIF1α) prolyl hydroxylases (PHDs) in cells ^4, 11, 33, 69–73^. To assess whether the increased MHC/HLA class I expression is associated with changes of ROS levels in the presence of vitC, we measured ROS levels along with MHC/HLA class I expression in cells in the presence of increasing concentrations of vitC. Notably, while vitC at a higher dose (2 mM) led to a significantly increased ROS level compared to the control (without vitC), lower doses of vitC (ranging from 0.1 to 1 mM) resulted in reduced ROS levels in these cells (**Figure S5C**), which is consistent with prior reports that vitC possesses both antioxidative and pro-oxidative properties, and at lower doses, it inhibits the formation of ROS in cells ^74–76^. Unlike the opposing effects of vitC on ROS at higher vs. lower concentrations, vitC induced MHC/HLA class I expression in a dose-dependent linear fashion from low to high concentrations (**Figure S5D**), suggesting that the induction of MHC/HLA class I expression is not mediated by an increase in ROS levels. To determine whether vitC induces MHC/HLA class I expression by acting as a co-factor for TET or PHD enzymes, we used Bobcat 339, a TET inhibitor ^77^, and IOX2, a PHD inhibitor ^78^, to inhibit their activities in cells. The TET inhibitor and PHD inhibitor were effective in reducing TET activity and increasing HIF1α levels in cells, respectively (**Figures S5E and S5F**). However, these inhibitors did not affect the vitC-induced increase in pSTAT1 and MHC/HLA class I expression (**Figure S5G**), suggesting that the increased levels of pSTAT1 and MHC/HLA class I expression induced by vitC are unlikely caused by the activation of TET and PHD enzymes in cells.

We next conducted co-culture experiments to explore the impact of vitC treatment on tumor cell interactions with immune cells. To assess whether vitC-pretreated PP tumor cells could activate antigen-presenting cells (APCs), we cocultured vitC-pretreated PP tumor cells with dendritic cells (DCs) derived from the bone marrow of naïve syngeneic FVB mice (**Figure 5D**). Notably, vitC- pretreated PP cells activated DCs, as evidenced by increased levels of MHC class II (MHC II), and co-stimulatory molecules CD86 and CD80 ^79, 80^ (**Figures 5E and S5H**). To further investigate the functional consequences of increased MHC/HLA class I expression on tumor cells induced by vitC, we employed ovalbumin (OVA)-expressing mouse tumor cell lines, B16-OVA and EL4-OVA ^68^. We confirmed that vitC increased pSTAT1 and MHC-I expression in both B16-OVA and EL4-OVA cells (**Figures 5F and S5I**). Co-culture of vitC-pretreated B16-OVA or EL4-OVA cells with MHC-I-restricted OVA-specific CD8^+^ T cells harvested from OT-I mice significantly increased CD8^+^ T cell proliferation (indicated by the negative CFSE population in CD8^+^ T cells) and production of anti-tumor cytokines, including IFNγ and tumor-necrosis factor-α (TNFα) ^68^ (**Figures 5G, 5H, S5J, and S5K**), indicating that vitC-pretreated tumor cells are able to stimulate T cell activation *in vitro*. Taken together, these results demonstrate that vitC treatment enhances the immunogenicity of tumor cells and their ability to activate immune cells.

### VitC induces vitcylation in tumor cells and enhances the STAT1-mediated immune responses *in vivo*

Finally, we examined how the tumor and tumor immune microenvironment respond to vitC treatment *in vivo*. We transplanted PP tumor cells into the mammary fat pads of syngeneic FVB mice and treated PP tumor-bearing mice with vitC (intraperitoneal injection, 4g/kg/day, 7 days), and tumors were harvested for analysis (**Figure 6A**). Consistent with our *in vitro* findings, we detected increased vitcylation and induced type I and II IFN responses in tumors upon vitC treatment (**Figures 6B and 6C**). Flow cytometry analysis showed that vitC treatment significantly increased pSTAT1 and MHC class I expression in tumor cells (**Figures 6D and 6E**). Additionally, we observed in the tumor microenvironment a significant increase in multiple immune cell populations, including DCs, CD4^+^, and CD8^+^ T cells, as well as enhanced activation of DCs and T cells (**Figures 6F-7H**). Similar results were obtained in E0771 tumor-bearing mice treated with vitC (**Figures S6A-S6F**). While we cannot exclude the direct effect of vitC on immune cells *in vivo*, our results suggest that vitcylation of tumor cells contributed, at least in part, to the activation of the immune milieu *in vivo*. Collectively, our *in vitro* and *in vivo* results suggest that STAT1 vitcylation induced by vitC enhances STAT1 phosphorylation and subsequently increases antigen processing and presentation in tumor cells, which in turn can activate multiple populations of immune cells, including DCs and T cells, in the tumor microenvironment (**Figure 7**).

**Figure 6.**
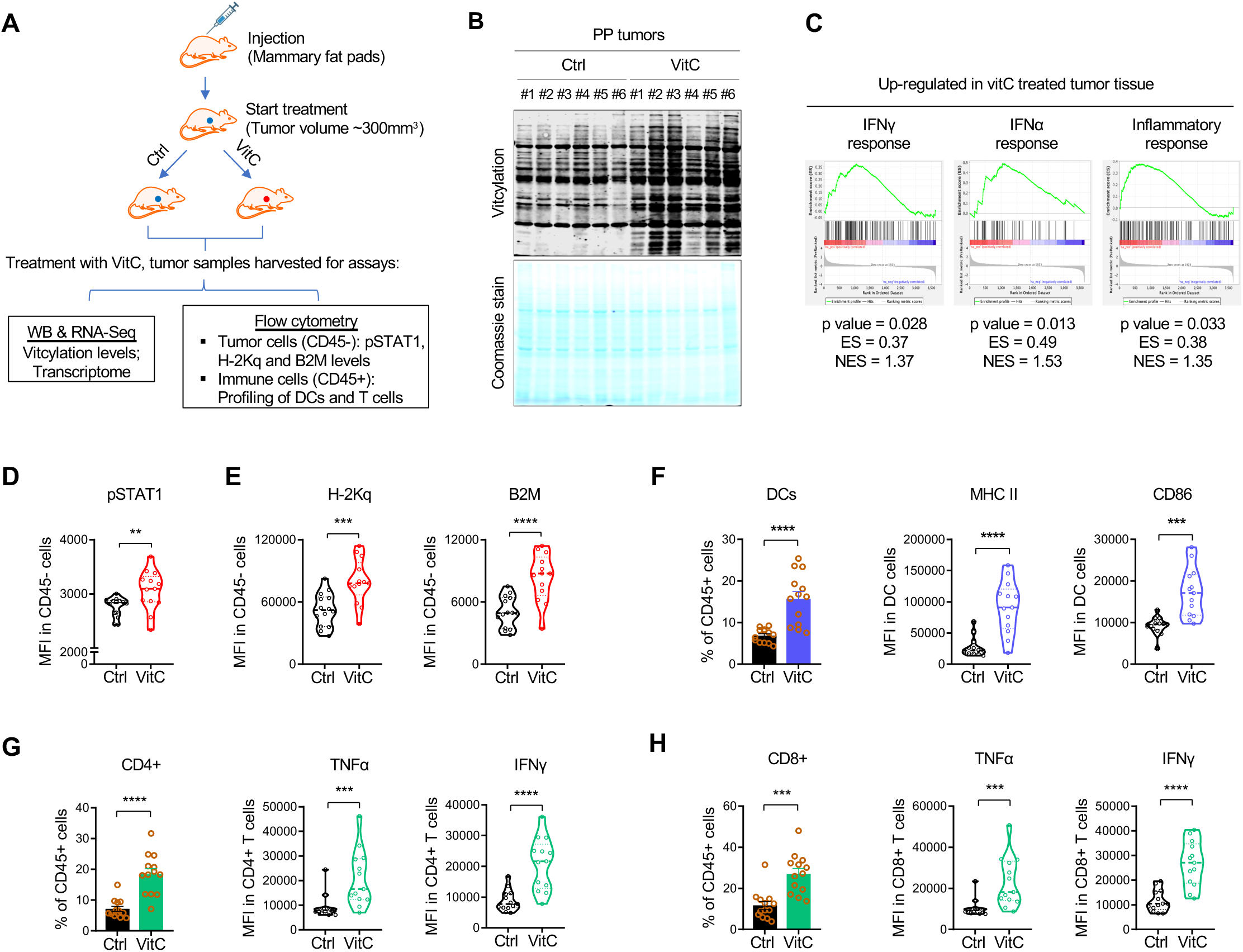
VitC induces vitcylation in tumor cells *in vivo* with increased STAT1-mediated immune responses. (A) Workflow for analyzing vitcylation level and immune cell infiltration in tumors *in vivo*. PP tumor cells were injected into the mammary fat pads of syngeneic females. Tumor-bearing mice were administered vitC (i.p. 3mg/kg, qd) when tumor volume reached approximately 300 mm^3^. After treatment, tumor tissues were harvested for analyses for vitcylation and immune response. (B) Vitcylation levels in PP tumors were measured by WB with anti-vitcylation antibody (n = 6 for each group). Protein levels were normalized by coomassie staining. (C) Top-ranked upregulated GSEA signatures in the tumor tissue of vitC-treated PP tumor-bearing mice at 7 days (n = 3). (D and E) Flow cytometry analysis of pSTAT1 (D), H-2Kq, and B2M (E) expression on PP tumor cells (CD45^-^) from vehicle- or vitC-treated mice (vehicle n = 14, vitC treated n = 13). Data are represented as mean ± SEM. **p < 0.01, ***p < 0.001, ****p < 0.0001. (F) Flow cytometry analysis of DCs (CD45^+^ CD11c^+^) population, and the MHC II and CD86 expression in DCs in PP tumor tissue from vehicle- or vitC-treated mice (vehicle n = 14, vitC treated n = 13). Data are represented as mean ± SEM. ***p < 0.001, ****p < 0.0001. (G) Flow cytometry analysis of CD4^+^, CD4^+^ TNFα^+^, and CD4^+^ IFNγ^+^ T cells (CD45^+^ CD3^+^) isolated from PP tumors tissue from vehicle- or vitC-treated mice (vehicle n = 14, vitC treated n = 13). Data are represented as mean ± SEM. ***p < 0.001, ****p < 0.0001. (H) Flow cytometry analysis of CD8^+^, CD8^+^ TNFα^+^ and CD8^+^ IFNγ^+^ T cells (CD45^+^ CD3^+^) isolated from PP tumors tissue from vehicle- or vitC-treated mice (vehicle n = 14, vitC treated n = 13). Data are represented as mean ± SEM. ***p < 0.001, ****p < 0.0001. See also Figure S6.

**Figure 7.**
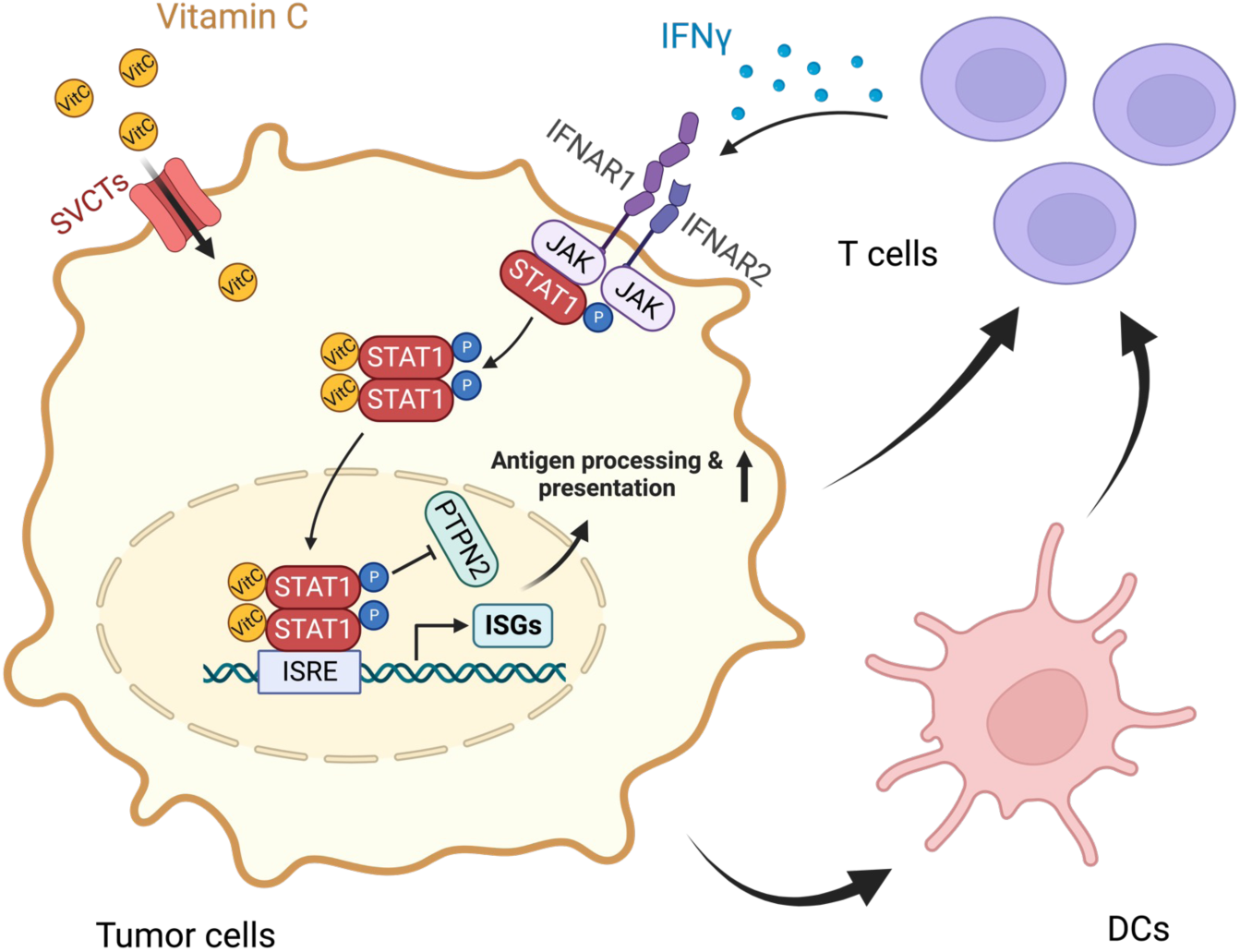
STAT1 vitcylation by vitC enhances STAT1-mediated IFN signaling pathway and immune response in tumor cells. VitC can be actively imported into cells through sodium-ascorbate co-transporters, also referred to as sodium-dependent vitamin C transporters (SVCTs). VitC vitcylates STAT1in cells, resulting in sustained phosphorylation and nuclear localization of STAT1 through attenuation of dephosphorylation by protein phosphatase PTPN2. This in turn upregulates STAT1-mediated gene expression and IFN pathway activation, leading to enhanced antigen processing and presentation and antitumor immunity in tumor cells.

## DISCUSSION

The understanding of vitC as a potential avenue for cancer therapy is still evolving. In this study, we show that vitC can perform enzyme-independent modifications on lysine in peptides and proteins through a process that we termed “vitcylation” to distinguish it from the previously described ascorbylation induced by DHA ^26–29^. In contrast to DHA-induced ascorbylation/glycation under acidic pH conditions, vitcylation occurs across a broader range of pH levels, ranging from 7 to 10, with the peak activity observed around pH 9-10. This suggests that vitC’s modification activity is optimal under alkaline conditions rather than acidic conditions. Furthermore, the pH and dose-dependent nature of this modification may provide evidence for the potential therapeutic efficacy of high-dose vitC and suggest that high-dose vitC may have specific benefits and effects that are distinct from lower doses. Understanding the pH and dose-dependent mechanisms of vitC’s activity adds to the body of knowledge surrounding this compound and its potential therapeutic applications such as cancer treatment.

By modifying lysine residues, vitcylation may regulate protein-protein interactions, enzymatic activity, or protein stability, thereby influencing the function of the modified proteins. The discovery of vitcylation as a form of PTM opens a new avenue for understanding the role of vitC in cellular processes and disease mechanisms. We described here the broad range of vitcylated proteins and their associated functions, including metabolic pathway regulation, DNA repair, signal transduction, ATPase activity, and telomere maintenance. Further identification and characterization of these vitcylated proteins will provide valuable insights into how vitC may contribute to the regulation of cell physiology and influencing multiple aspects of cellular functions.

Furthermore, lysine vitcylation has the potential to serve as a tangible biomarker for various physiological and pathological conditions. By studying the specific vitcylated proteins and their modification patterns, researchers can potentially identify unique signatures or patterns associated with certain diseases or physiological states. These vitcylation markers could be used to develop diagnostic tools or biomarker panels for assessing the status of certain conditions or monitoring treatment responses. This information can potentially guide treatment decisions and improve patient outcomes, especially in cancer treatment, since a major challenge in studying high-dose vitC as a cancer therapy is the determination of appropriate vitC dosage and developing reliable biomarkers to monitor effects.

Going beyond the identification of vitcylation, we further found that vitcylation of STAT1-K298 has significant implications for the conformational transition and subsequent function of STAT1. A Rosetta molecular modeling analysis revealed that this modification interferes with the transition of pSTAT1 from a parallel conformation to an anti-parallel conformation, which is crucial for the binding of PTPN2 and the subsequent dephosphorylation of pSTAT1. As a result, this altered interaction prevents dephosphorylation of pSTAT1, leading to elevated pSTAT1 levels and enhanced STAT1-mediated IFN response pathway and immune responses.

Interestingly, the genetic mutation STAT1-K298N has been identified as a causative factor in certain autoimmune diseases. This mutation involves the substitution of lysine (K) with asparagine (N) at position 298 within the STAT1 protein. Our Rosetta molecular modeling analysis also predicted that this alteration affects the ability of pSTAT1 to transition from a parallel conformation to an anti-parallel conformation, leading to compromised binding of PTPN2 for dephosphorylation. This is consistent with the previous report that STAT1-K298N has impaired binding to PTPN2 and increased pSTAT1 ^55^. The shared underlying molecular mechanism between STAT1-K298N and STAT1-vitcyl-K298 suggests a critical role for this specific lysine residue in the regulation of STAT1 signaling and immune responses. However, it is important to differentiate between the mutation and the modification, as they have opposite effects on immune regulation. While the STAT1-K298N mutation can lead to chronic upregulation of immune responses, contributing to autoimmune diseases, the STAT1-K298-vitcylation, by contrast, can augment immune function in a manner that is beneficial for combating pathological conditions, such as cancer and viral infections.

Finally, lysine vitcylation not only provide a unique window into the understanding the role of vitC in protein modification and cellular regulation, but also expands our broader knowledge of protein modifications and their contributions to health and disease. It highlights the significance of vitC as a regulatory factor and underscores the complex interplay between nutrients, cellular processes, and physiology. Further research will help elucidate the full extent of the vitcylation and its relevance in various biological processes, including potential implications for cancer treatment and other therapeutic interventions.

## ACKNOWLEDGEMENTS

We thank Drs. Marie Bao, Thomas Roberts, and Bingqiu Xiu for scientific discussion. This work was supported in part by grants from Department of Defense Breast Cancer Research Program Breakthrough Award HT9425-23-1-0026 (Q.W.), Breast Cancer Research Foundation (J.J.Z.), and National Health Institute (NIH) P50 CA168504 (J.J.Z) and CA210057 (J.J.Z).

## AUTHOR CONTRIBUTIONS

X.H. and J.J.Z conceived and designed the study and wrote the manuscript. X.H. performed most of the experiments. Y. W. and J.W. helped with structure analysis; T.J. and W.W. helped with *in vivo* treatments; S. X. conducted transcriptomic assays; J.M.A. and J.L. performed mass spectrometric analysis; Q.W. and H.G. helped with flow cytometry; J.Z. and S.L. performed the thermostability change calculation. J.S.B. and Q.W. provided critical materials; Y.W., J.W., Q.W., J.S.B., K.Z., and L.C.C. contributed to scientific discussions. X.H., Y.W., Q.W., L.C.C., and J.J.Z. reviewed and edited the manuscript.

## STAR ★ METHODS

### RESOURCE AVAILABILITY

#### Lead contact

Further information and requests for resources and reagents should be directed to and will be fulfilled by the lead contact, Jean J. Zhao

#### Materials availability

Plasmids generated in this study are available from the lead contact upon request.

### EXPERIMENTAL MODEL AND SUBJECT DETAILS

#### Cell lines

Cells were cultured under standard conditions in a humidified incubator with 5% CO_2_ at 37℃. Cal-51, MCF7, E0771, PC-9, HCT116, and HeLa cells were obtained from the American Type Culture Collection (ATCC), verified to be negative for mycoplasma, and authenticated by short tandem repeat analysis using the Promega GenePrint 10 System. Cal-51, MCF7, PC-9, HCT116, and HeLa cells were cultured in RPMI 1640 (Gibco) supplemented with 10% fetal bovine serum (FBS) (Life Technologies) and PenStrep (Hyclone). E0771 cells were cultured in RPMI 1640 supplemented with 10% FBS, PenStrep, and 10 mM HEPES (Life Technologies, 15630080). B16- OVA and EL4-OVA cells were cultured in DMEM (Gibco) supplemented with 10% FBS and 250 μg/ml G418 (Invitrogen). PP cells were derived from mouse mammary tumors and cultured in DMEM/F12 media supplemented with 10% FBS, 25 ng/ml hydrocortisone, 5 μg/ml insulin, 8.5 ng/ml cholera toxin, 0.125 ng/ml epidermal growth factor (EGF), 5 μM Y-27632 Rock1 inhibitor, and PenStrep, as previously described ^81^.

#### Animals

All mouse experiments were conducted in compliance with federal laws and institutional guidelines, as approved by the Institutional Animal Care and Use Committees of Dana-Farber Cancer Institute and Harvard Medical School. The relevant animal protocols limited the maximum tumor diameter to 25 mm, which was not exceeded in any experiment. CO_2_ inhalation was used to euthanize the mice. Female wild-type FVB and C57BL/6 mice, aged six-to-eight weeks, were obtained from the Jackson Laboratory. For tumor formation assays, 5×10^5^ PP cells or 10^5^ E0771 cells were injected into the thorracic fat pad in 50% matrigel. VitC (Sigma-Aldrich, A4034) was prepared daily by resuspending the powder in PBS (Hyclone). Intraperitoneal administration of vitC was conducted at a dose of 4 g/kg, qd. Mice in the control group were treated with PBS.

### METHOD DETAILS

#### Transcriptome methodology

An Ion AmpliSeq Custom Panel containing 4,604 cancer- and immune-associated genes (designed by Thermo Fisher using Ion AmpliSeq designer) was utilized for our studies, as previously described ^68^. For each sample, 10 ng total RNA was used to prepare the cDNA library. Libraries were multiplexed and amplified using an Ion OneTouch 2 System and sequenced on an Ion Torrent Proton system (Thermo Fisher). Count data was generated using Thermo Fisher’s Torrent suite and ampliSeqRNA analysis plugin. For Gene Ontology enrichment and KEGG pathway analysis, genes with a mean fold change (vitC treated vs control) greater than 2 or lesser than 0.5 were utilized. Gene Ontology enrichment and KEGG pathway analysis were carried out using Cytoscape Software and STRING plugin. For GSEA analysis, genes were first ranked according to log_2_(fold change) and then analyzed using the GSEAPreanked tool with MsigDB v7.1 Hallmarks gene sets and the ’classic’ method ^83^.

#### Bioinformatic analysis

The subcellular localization of vitcylated proteins was predicted using WoLF PSORT, a subcellular localization prediction program (https://wolfpsort.hgc.jp) ^84^. Gene Ontology enrichment analysis for the vitcylated proteins was carried out using DAVID bioinformatics resources 6.8 (https://david.ncifcrf.gov/home.jsp) ^85^. KEGG pathway analysis was performed using KOBAS (http://kobas.cbi.pku.edu.cn/genelist/) ^86^. For vitcylation motif analysis, the 10 amino acid residues (−10 to +10) on either side of the vitcylation site were selected and a consensus logo was generated using the WebLogo webserver (http://weblogo.berkeley.edu/logo.cgi) ^87^.

#### Flow cytometry

For tumor cell lines, one million cells were stained with the appropriate antibodies diluted in PBS (Hyclone) plus 2% FBS (Life Technologies) for 30 min on ice. For tumor samples, tumors were mechanically disrupted by chopping and chemically digested in dissociation buffer (2 mg/ml collagenase type IV (Worthington Biochemical), 0.02 mg/ml DNase (Sigma Aldrich) in DMEM (Life Technologies) containing 5% FBS (Life Technologies), PenStrep (Hyclone) with agitation at 37℃ for 45 min. After lysing of red blood cells (BD Biosciences), single-cell suspensions were incubated with appropriate antibodies for 30 min on ice.

For human antibodies, antibodies were purchased from BioLegend unless otherwise indicated: pSTAT1(Y701) (clone A17012A), HLA (clone W6/32), B2M (clone A17082A). Mouse antibodies: pSTAT1(Y701) (Cell Signaling Technology, 8062S), H-2Kq (clone KH114), H-2Kb (clone 28-8- 6), B2M (clone A16041A), CD45 (clone 30-F11), CD3 (clone 145-2C11), CD4 (clone RM4-5), CD8 (clone HIT8a), IFNγ (clone XMG12), TNFα (clone MP6-XT22), CD11c (clone N418), CD86 (clone GL-1), CD80 (clone 16-10A1), MHC II (clone M5/114.15.2), CD103 (clone 2E7). Anti-mouse/rat FoxP3 staining set (eBioscience) was used for intracellular staining according to the manufacturer‘s instructions. For IFNγ and TNFα analysis, cells were stimulated *in vitro* with the Leukocyte Activation Cocktail with protein transport inhibitor Brefeldin A (BD Biosciences, 550583).

#### Western blots

Western blotting was performed as previously described ^68^ using the following antibodies: anti- vitcylation antibody (generated by Abclonal Technology) and Cell Signaling Technology antibodies to STAT1 (14994S), GAPDH (5174S), ACTIN (4967S), GFP (2955S), HA (2367S), and COX4 (4850S).

#### Dot-blot assays

Peptides were spotted on nitrocellulose membranes. After the membrane dried, the membrane was blocked with 5% skimmed milk in TBST for 1 hour, followed by the incubation with the anti- vitcylation antibody overnight at 4℃ and the secondary antibody overnight at 4℃. After washing three times with TBST, the membrane was scanned by Odyssey Dlx Imaging System (LI-COR).

#### Thermostability change calculation

The antiparallel dimer structure from PDB 1YVL (Asymmetric Unit) is initially considered with 3.00Å X-ray resolution. The whole calculation includes several steps as below:

1. The input structure preparation

The wild type K298, the mutation K298N and the vitcyl-K298 (in two chains) were relaxed with Cartesian coordination type and constraints ^88–90^. The option/flag was set as follows.

-nstruct 200

-ex1

-ex2

-relax:constrain_relax_to_start_coords

-ramp_constraints true

-use_input_sc

-flip_HNQ

-no_optH false

-relax:cartesian

-beta_nov16_cart

-corrections::beta_nov16

-crystal_refine

-in:auto_setup_metals

-extra_res PTM.params

The structures with the lowest scores were used in the next step.

2. The preparation of params and rotlib files

To make the vitcyl-K298 recognized by Rosetta, the params and rotlib files were generated by related modules using Lysine as reference ^91, 92^.

3. MD sampling

The side chain of vitcyl-K298 has more than 10 chi angles, which is beyond the recommended limitation of Rosetta. To better find the low energy conformations of vitcyl-K298, we adopted molecular dynamics (MD) simulations to perform the sampling. Ff14SB was used to parameterize the protein ^93^. The vitcyl-K298 was parameterized using GAFF and its partial charge was parameterized using RESP ^94, 95^.

10ns MD simulations were performed using Amber20 and 100 frames of snapshots were extracted from the last 5ns scoring using Rosetta. A 100kcal/mol positional restraints were given on residues 9 Å away from α-carbon of K298 to minimize the noise arose from the thermal fluctuation during the MD simulations.

4. Scoring and ddG calculation

The score_jd2 module was used with the following options ^96, 97^.

-beta_nov16_cart

-corrections::beta_nov16

-fa_max_dis 9.0

-extra_res VLY.params

Average scores for K298, N298 and vitcyl-K298 were calculated on the 100 frames of snapshots. Values of ddG were calculated from the difference as:

ddG = Score avg(Mut/Mod) – Score ave(WT)

#### Cellular levels of vitC, ROS, and TET activity assay

Cellular vitC assay kit (Cayman, 700420), ROS/superoxide detection assay kit (abcam, ab139476), and epigenase 5mC-hydroxylase TET activity/inhibition assay kit (Epigentek, P-3086-48) were used for cellular vitC level assay, ROS level assay, and TET activity assay according to the manufacturer’s instructions, respectively.

#### RT-PCR

RT-PCR was performed as previously described ^98^. Primer sequences used for RT-PCR were as follows. *Tap1* (mouse) forward: 5’- GGACTTGCCTTGTTCCGAGAG-3’; reverse: 5’- GCTGCCACATAACTGATAGCGA-3’. *Lmp2* (mouse) forward: 5’- ATGTGGTACTCAATTCACAAGCA-3’; reverse: 5’-AAGCAAGGATGGTTCCTGGAG-3’. *B2m* (mouse) forward: 5’-TTCTGGTGCTTGTCTCACTGA-3’; reverse: 5’- CAGTATGTTCGGCTTCCCATTC-3’. *Irf1* (mouse) forward: 5’- GTTGTGCCATGAACTCCCTG-3’; reverse: 5’-GTGTCCGGGCTAACATCTCC-3’. *H2k1* (mouse) forward: 5’- CAGGTGGAGCCCGAGTATTG-3’; reverse: 5’- CGTACATCCGTTGGAACGTG-3’. *Actin* (mouse) forward: 5’- CGCCACCAGTTCGCCATGGA-3’; reverse: 5’- TACAGCCCGGGGAGCATCGT-3’. *HLA-B* (human) forward: 5’- CAGTTCGTGAGGTTCGACAG-3’; reverse: 5’- CAGCCGTACATGCTCTGGA-3’. *TAP1* (human) forward: 5’-CTGGGGAAGTCACCCTACC-3’; reverse: 5’- CAGAGGCTCCCGAGTTTGTG-3’. *TAP2* (human) forward: 5’- TGGACGCGGCTTTACTGTG-3’; reverse: 5’- GCAGCCCTCTTAGCTTTAGCA-3’. *LMP2* (human) forward: 5’- GCACCAACCGGGGACTTAC-3’; reverse: 5’- CACTCGGGAATCAGAACCCAT-3’. *B2M* (human) forward: 5’- GAGGCTATCCAGCGTACTCCA-3’; reverse: 5’- CGGCAGGCATACTCATCTTTT-3’. *ACTIN* (human) forward: 5’-CACCAACTGGGACGACAT-3’; reverse: 5’- ACAGCCTGGATAGCAACG-3’. Relative copy number was determined by calculating the fold change difference in the gene of interest relative to *Actin* (mouse) or *ACTIN* (human). RT-PCR was performed on an Applied Biosystems 7300 machine.

#### Generation of mouse DCs

Mouse DCs were obtained from the bone marrow of FVB/NJ mice by modifying the previously described protocol ^99^. For DC generation, bone marrow cells were cultured in RPMI 1640 supplemented with 20 ng/ml GM-CSF (Stem Cell Technologies, 78017), 10% FBS, and 100 μg/ml PenStrep. Fresh RPMI 1640 with 20 ng/ml GM-CSF, 10% FBS, and 100 μg/ml PenStrep was added after 3 days, and non-adherent cells (DCs) were harvested and co-cultured with PP cells after another 2 days.

#### Co-culture experiments

For *in vitro* co-culture of tumor cells with CD8^+^ T cells, B16-OVA and EL4-OVA cells were pretreated with vitC or PBS for 3 days. CD8^+^ T cells were isolated from spleens of OT-I mice using a CD8a^+^ T-cell isolation kit (StemCell Technologies) with an autoMACS Pro Separator. Isolated CD8^+^ T cells were suspended in RPMI 1640 medium (Gibco) with 5% FBS, labeled with 5 μM CFSE (Biolegend) for 10 min in the dark at room temperature, and washed twice in 10× volume of T cell media (RPMI 1640 with 10% FBS and 55 μM 2-mercaptoethanol (Gibco)). One hundred thousand CD8^+^ T cells were co-cultured with vitC- or control-pretreated tumor cells at a ratio of 1:8 tumor cells: T cells in RPMI 1640 supplemented with CD3/CD28 Dynabeads (1:1 ratio of cells: beads, ThermoFisher) 2.5 ng/ml IL-7 (Biolegend), 50 ng/ml IL-15 (Biolegend), and 2 ng/ml IL-2 (Biolegend) for 2 days at 37℃ in the dark. At the experimental endpoint, CD8^+^ T cell proliferation and activation were analyzed by flow cytometry.

For *in vitro* co-culture of tumor cells with DCs, PP cells were pretreated with vitC or PBS for 3 days. One hundred thousand mouse bone marrow-derived DCs were co-cultured with vitC- or control-pretreated PP cells at a ratio of 1:4 tumor cells: DCs in RPMI 1640 supplemented with 20 ng/ml GM-CSF, 10% FBS and lipofectamine 3000 (2 μl/ml, Invitrogen) for 2 days at 37℃. At the experimental endpoint, DCs activation was analyzed by flow cytometry.

#### Generation of STAT1-deficient cells

CRISPR-Cas9 genome editing systems were used to generate STAT1-deficient cells ^100^. Oligos sequences for guide RNAs were as follows. *Stat1* (mouse) #1 forward: 5’- CACCGGAACCCCCCGTGCGCGTGG-3’; #1 reverse: 5’-AAACCCACGCGCACGGGGGGTTCC-3’. *Stat1* (mouse) #2 forward: 5’-CACCGGTCGCAAACGAGACATCAT-3’; #2 reverse: 5’-AAACATGATGTCTCGTTTGCGACC-3’. Annealed guide oligos were cloned into the CRISPR- Cas9 expression vector PX458. The constructed sg*Stat1*(mouse)-PX458 plasmids were transfected into PP cells. The next day, single GFP^+^ cells were sorted into 96-well plates by flow cytometry. Two weeks later, colonies emerged and single colonies were expanded into 6 well plates. STAT1 knockout cells and control cells were selected by western blot.

#### Co-immunoprecipitation (Co-IP) and immunoprecipitation (IP)

Cells were lysed using lysis buffer (50 mM Tris-HCl (pH 7.4), 150 mM NaCl and 0.1% NP-40 (for Co-IP) or 0.5% NP-40 (for IP)) at 4℃ for 30 minutes. Cell lysates were then centrifuged at 12,000 rcf for 15 min and the supernatants were collected and incubated with GFP-Trap magnetic agarose (Chromotek, gtma-20) overnight at 4℃. After washing the beads with lysis buffer three times, they were resuspended in 1× western blotting loading buffer and denatured at 95℃ for 10 minutes.

#### Immunofluorescence

For endogenous STAT1 immunofluorescence, cultured cells were fixed with 4% formaldehyde in PBS for 15 min at room temperature. The fixative was aspirated, and the cells were rinsed three times in PBS for 5 min each. The cells were then covered with ice-cold 100% methanol and incubated for 10 min at −20℃. After incubation, the cells were rinsed in PBS for 5 min. The specimen was blocked in blocking buffer (5% normal serum from the same species as the secondary antibody and 0.3% Triton X-100 in PBS) for 60 min. The blocking solution was aspirated, and diluted STAT1 antibody (Cell Signaling Technology, 14994) was applied and incubated overnight at 4℃. The cells were rinsed three times in PBS for 5 min each, followed by incubation with a fluorochrome-conjugated secondary antibody (ThermoFisher, A-11008) diluted in antibody dilution buffer (1% BSA and 0.3% Triton X-100 in PBS) for 2 hours at room temperature in the dark. The cells were then rinsed in PBS and incubated in PBS with DAPI (Cell Signaling Technology, 8961). Specimens were examined immediately using the appropriate excitation wavelength. For overexpressed STAT1-GFP immunofluorescence, cultured cells were fixed with 4% formaldehyde in PBS for 15 min at room temperature. The fixative was aspirated, and the cells were rinsed three times in PBS for 5 min each. The cells were then incubated in PBS with DAPI (Cell Signaling Technology, 8961). Specimens were examined immediately using the appropriate excitation wavelength.

### QUANTIFICATION AND STATISTICAL ANALYSIS

Statistical analysis was performed with Prism 9 (Graphpad Software Inc.). Two-way ANOVA with Tukey’s multiple comparisons test was used for tumor growth analysis. For other analysis, unpaired two-tailed Student’s test (for normally distributed data) and Mann-Whitney nonparametric test (for skewed data that deviated from normality) were used to compare two conditions. One-way ANOVA with Tukey’s multiple comparisons test (for normally distributed data) and Kruskal-Wallis nonparametric test (for skewed data) were used to compare three or more means. Quantitative data are expressed as means ± SEM. Differences with P < 0.05 were considered statistically significant; ns, not significant; * P < 0.05; ** P < 0.01; *** P < 0.001; **** P < 0.0001.

## SUPPLEMENTAL FIGURE LEGENDS

**Figure. S1.**
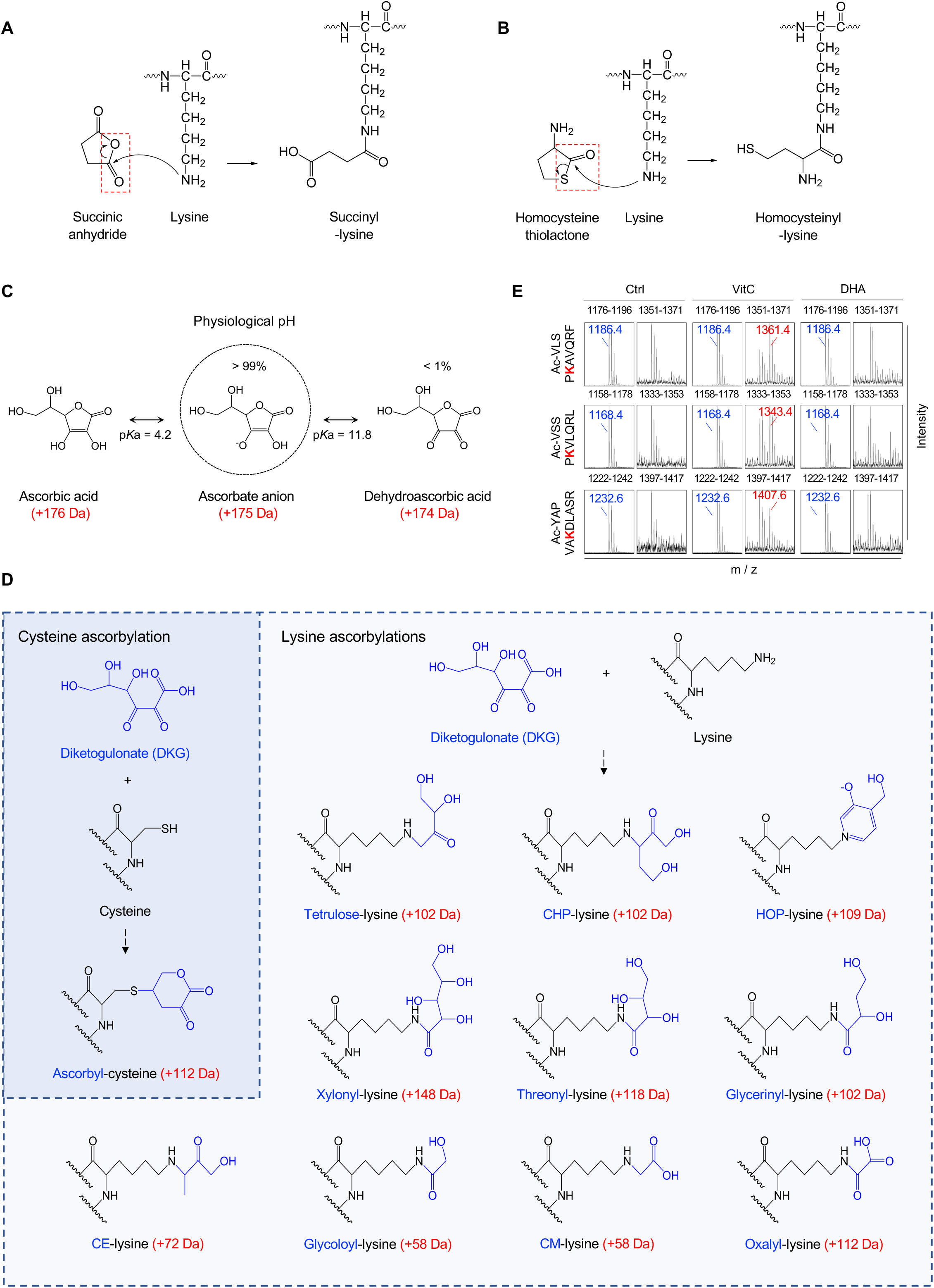
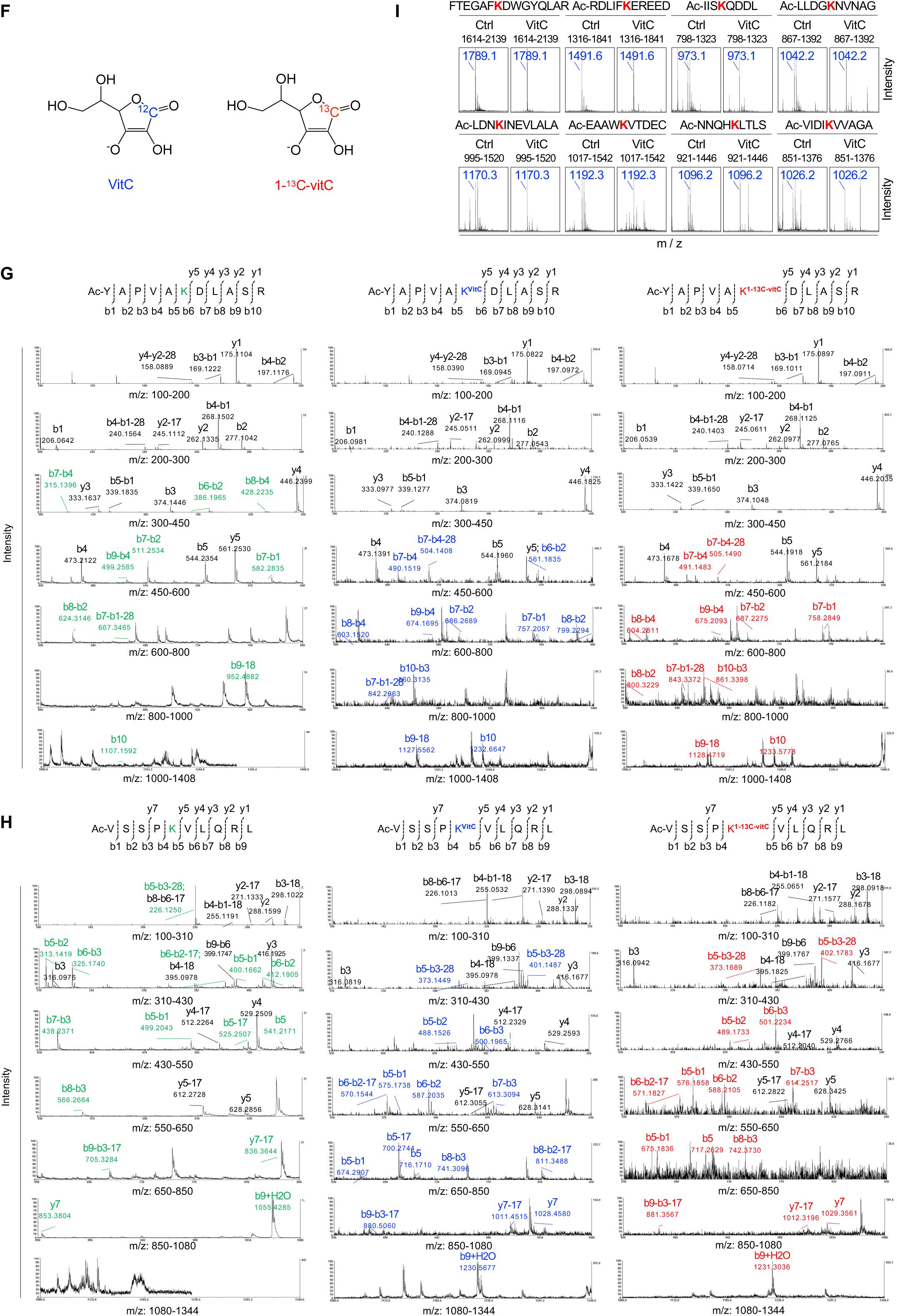
Formation of vitC-derived lysine vitcylation *in vitro*, related to figure 1. (A and B) Molecular mechanisms of lysine succinylation (A) and homocysteinylation formation (B) are presented, with the reactive lactone bonds circled for clarity. (C) The structures, molecular weights, and pKa’s of ascorbic acid, ascorbate anion, and dehydroascorbic acid are shown. (D) The structures of cysteine ascorbylation and lysine ascorbylations were illustrated to differentiate them from lysine vitcylation. (E) Synthetic lysine-containing peptides were incubated with vehicle, 2 mM vitC or 2 mM DHA at 37℃ for 3 hours, and the formation of vitcylated peptides was detected by MALDI-TOF/TOF MS. (F) The structures of vitC (left) and 1-^13^C-vitC (right), which were used in this study, are displayed. (G and H) MALDI-TOF/TOF MS/MS spectra of the unmodified peptide, vitcylated peptide, and 1-^13^C-vitcylated peptide (Ac-YAPVAKDLASR (G) and Ac-VSSPKVLQRL (H)) are shown, with lysine-containing unmodified fragments, vitcylated fragments, and 1-^13^C-vitcylated fragments marked by green, blue, and red colors, respectively. (I) Synthetic lysine-containing peptides (peptide sequences were listed above the spectrum) were incubated with vehicle or 2 mM vitC at 37℃ for 3 hours. The formation of vitcylated peptides was detected by MALDI-TOF/TOF MS.

**Figure. S2.**
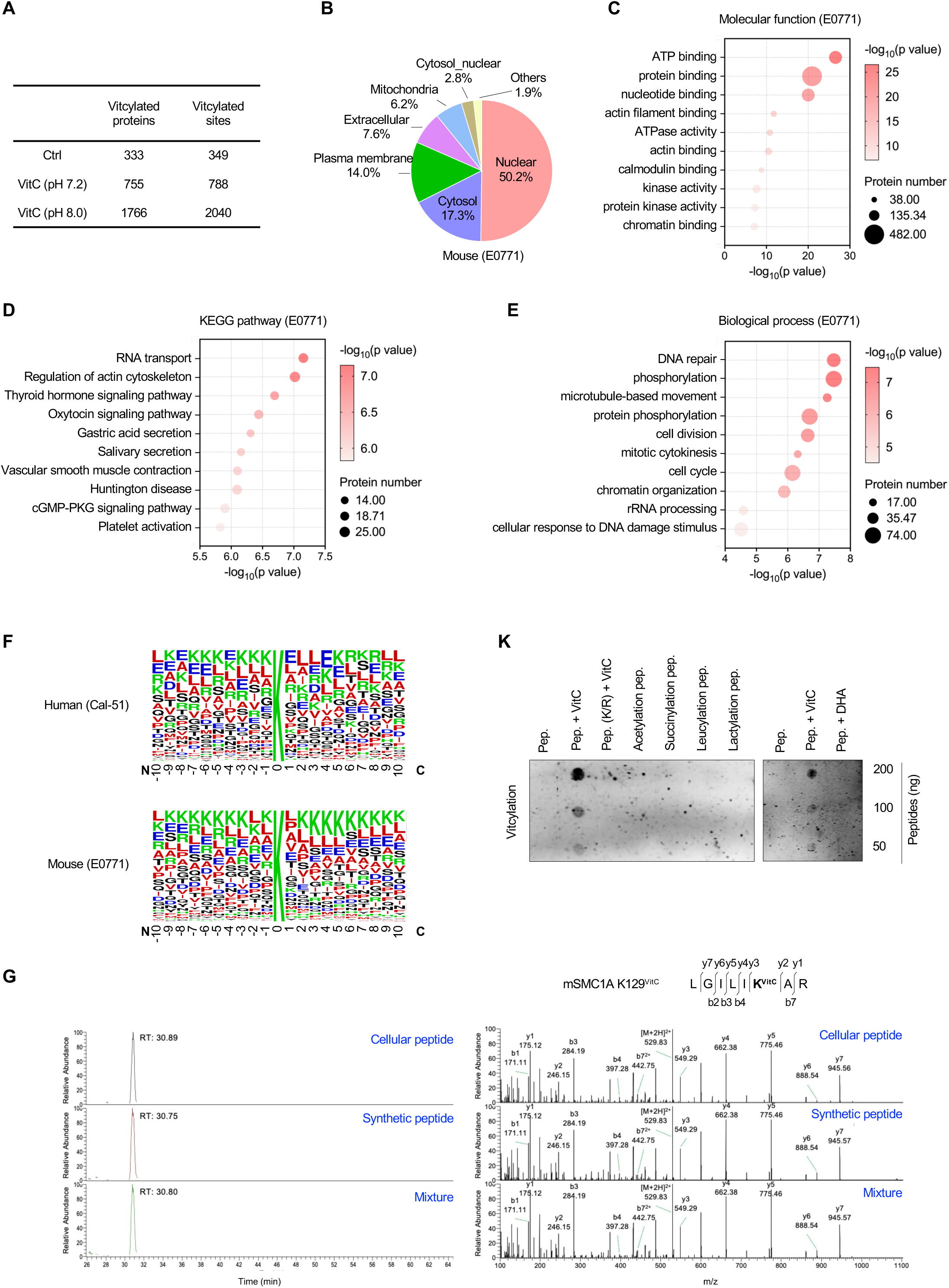
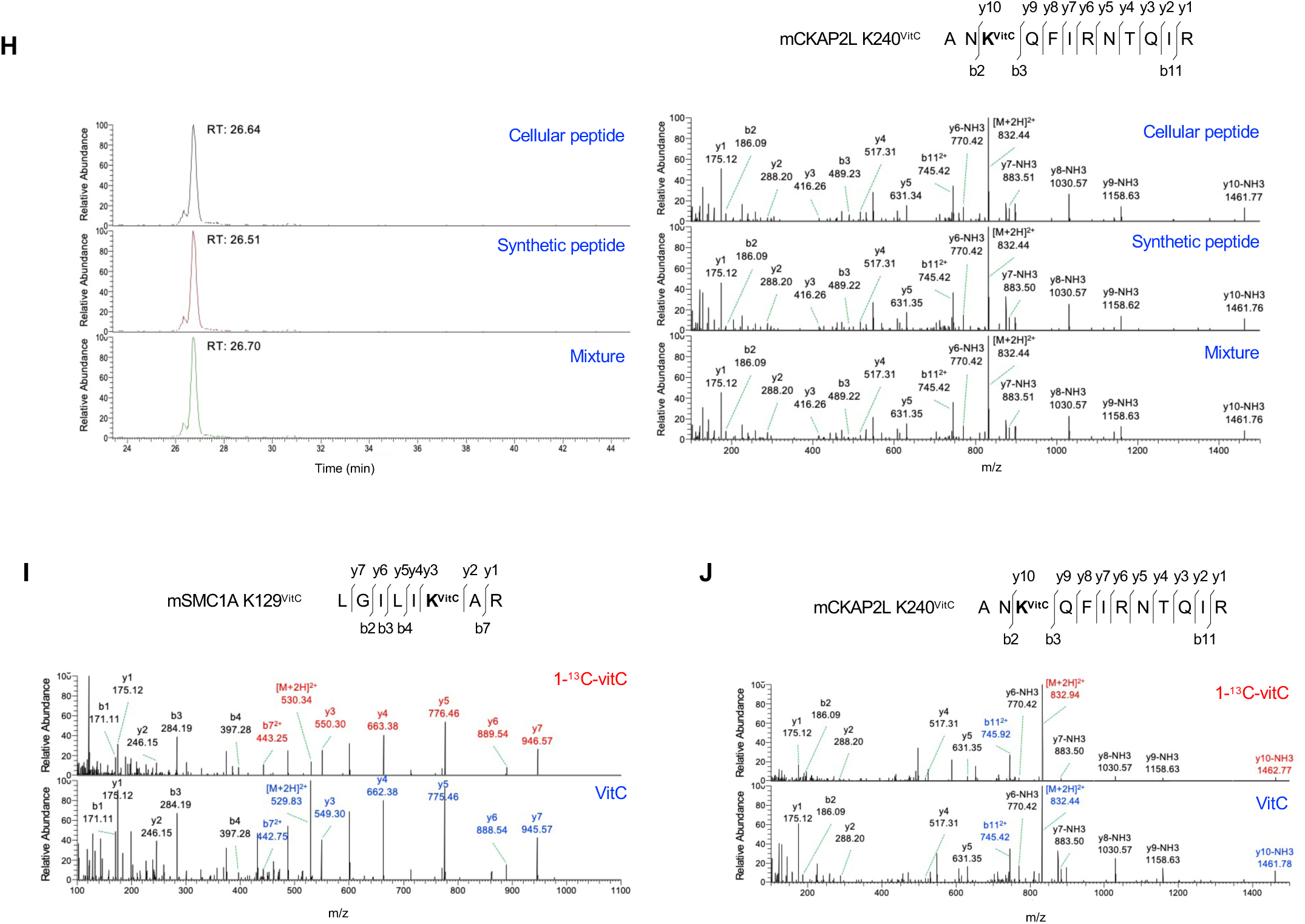
Lysine vitcylation exists in cells, related to figure 2. (A) Numbers of vitcylated proteins and sites identified in E0771 proteomic and E0771 proteomic incubated with vitC (1 mM for 12 hours) in indicated pH Tris-HCl buffer *in vitro* are summarized. E0771 proteomic was extracted by acetone. (B) Subcellular locations of lysine vitcylated proteins identified in E0771 (mouse) cells. The locations are classified as nuclear, cytosol, plasma membrane, extracellular, mitochondrial, cytosol_nuclear, and other compartments. (C) Top ten gene ontology molecular function enrichment of vitcylationo proteins identified in E0771 cells (mouse). (D) Top ten KEGG-based enrichment of lysine vitcylation proteins identified in E0771 cells (mouse). (E) Top ten gene ontology biological process enrichment of vitcylation proteins identified in E0771 cells (mouse). (F) Sequence probability logos of significantly enriched vitcylation site motifs for ±10 amino acids around the lysine vitcylation sites identified in Cal-51 cells (human) and E0771 cells (mouse). The size of each letter represents the frequency of the amino acid residue at that position. (G and H) Extracted ion chromatograms (left) and MS/MS spectra (right) from HPLC-MS/MS analysis of vitcylated peptides (mouse SMC1A, K129 (G) and mouse CKAP2L, K240 (H)) derived from E0771 (cellular peptide) respectively, their *in vitro* generated counterparts (synthetic peptide), and their mixture. (I and J) Extracted MS/MS spectra from HPLC-MS/MS analysis of 1-^13^C-vitcylated peptides and vitcylated peptides (mouse SMC1A, K129 (I) and mouse CKAP2L, K240 (J)) derived from E0771 cells (in cells, the lysine-containing 1-^13^C-vitcylated fragments and vitcylated fragments were marked by red and blue colors, respectively). (K) The pan anti-vitcylation antibody was tested for its reactivity with vitcylated peptide (pep. + vitC, peptide pre-incubated with 2 mM vitC at 37℃ for 3 hours) and cross-reactivity with unmodified (pep. and pep. (K/R) + vitC), DHA-derived modification peptide (pep. + DHA, peptide pre-incubated with 2 mM DHA at 37℃ for 3 hours), as well as synthetic acetylated, succinylated, leucylated and lactylated peptides (peptide sequence: Ac-VLSPKAVQRF).

**Figure. S3.**
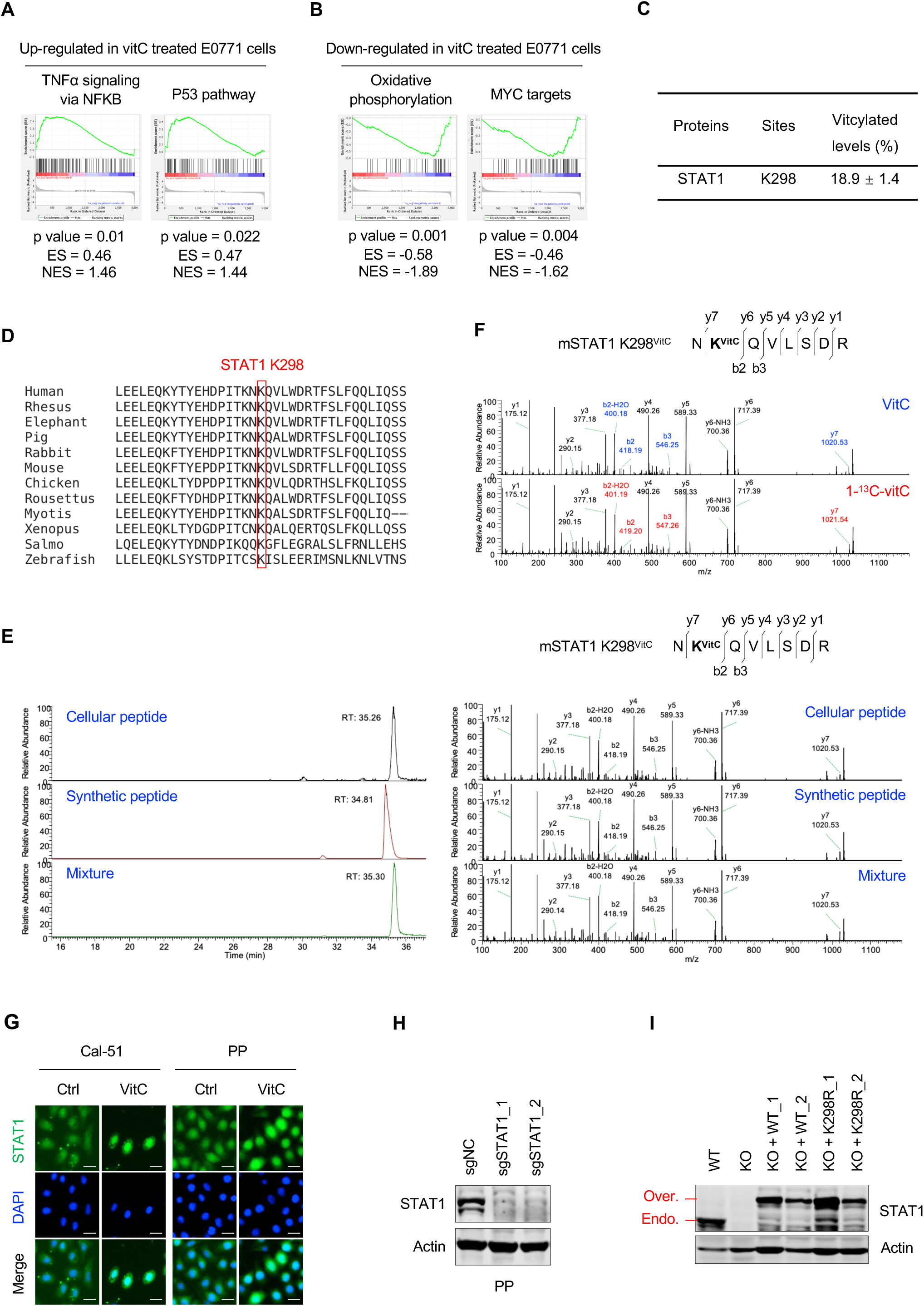
Vitcylation of STAT1 K298 activates STAT1, related to Figure 3. (A and B) Upregulated (A) and downregulated (B) GSEA signatures were observed in E0771 cells treated with 2 mM vitC for 2 days (n = 3). (C) Levels of vitcylation of human STAT1 K298 were determined by HPLC-MS/MS analysis. Data are represented as mean ± SEM. (D) Sequence analysis of STAT1 K298 site from multiple vertebrate species. (E) Extracted ion chromatograms (left) and MS/MS spectra (right) from HPLC-MS/MS analysis of a vitcylated peptide (mouse STAT1, K298) derived from E0771 (cellular peptide), its *in vitro* generated counterparts (synthetic peptide), and their mixture. (F) Extracted MS/MS spectra from HPLC-MS/MS analysis of vitcylated peptides (upper) and 1- ^13^C-vitcylated peptides (lower) (mouse STAT1, K298) derived from E0771 cells. The lysine-containing vitcylated fragments and 1-^13^C-vitcylated fragments were marked by blue and red colors, respectively). (G) Nuclear translocation of endogenous STAT1 in Cal-51 and PP cells cultured in vehicle- or vitC-containing medium for 2 days was assessed by immunofluorescence (0.5 mM vitC for Cal- 51 culture, 2 mM vitC for PP culture). Green represents anti-STAT1 and blue represents DAPI, with merged images allowing assessment of nuclear localization of STAT1, hereafter referred to as STAT1 immunofluorescence (scale bar, 50 μM). (H) Western blots for STAT1 and Actin in PP-sg_NC, PP-sgSTAT1_1 and PP-sgSTAT1_2 cells. (I) Western blots for STAT1 and Actin in PP-sgNC, PP-sgSTAT1_1 and PP-sgSTAT1 cells re-expressing STAT1-WT-GFP or re-expressing STAT1-K298R-GFP.

**Figure. S4.**
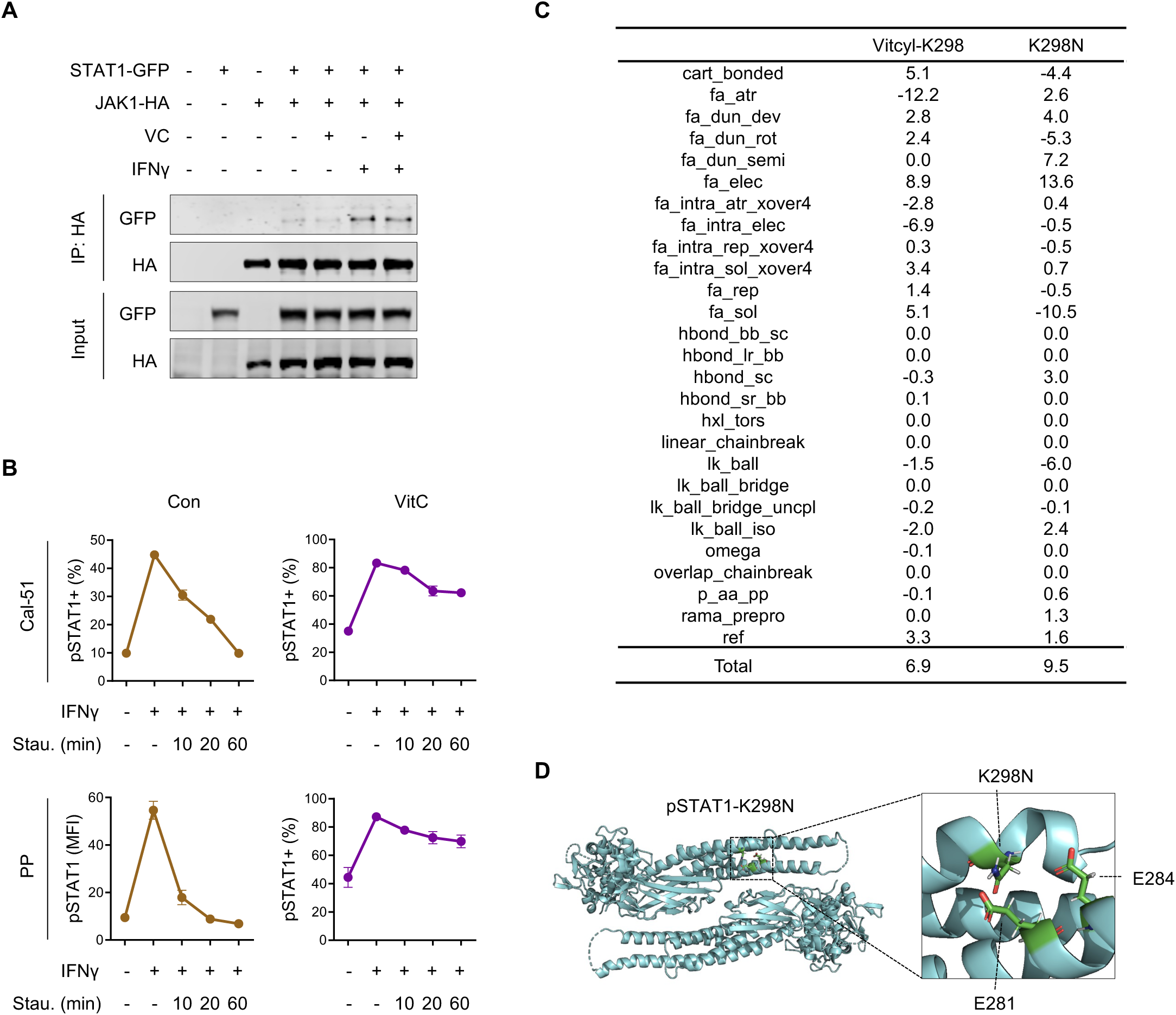
STAT1 K298 vitcylation impairs STAT1 dephosphorylation, related to Figure 4. (A) HeLa cells co-expressing STAT-GFP and JAK1-HA were treated with vehicle or 300 μM vitC for 1 day, followed by stimulation with 100 ng/ml IFNγ for 15 min. Interaction between STAT1 and JAK1 was assayed by co-immunoprecipitation. (B) Cells were pretreated with vehicle or vitC (0.2 mM vitC for Cal-51 cell culture, 1 mM vitC for PP cell culture) for 2 days, then cells were stimulated with 100 ng/ml IFNγ for 15 min followed by incubation with 1 μM staurosporine for indicated times. The pSTAT1 levels were measured by flow cytometry immediately (n = 3). Data are represented as mean ± SEM. (C) Energy contribution difference of STAT1 K298N and K298 vitcylation. Numbers are reported in Rosetta energy unit. (D) Structures of pSTAT1 with K298N mutation in the antiparallel dimer conformation from the last snapshot of MD simulation. STAT1 K298N lose the salt bridges of K298/E281 and K298/E284.

**Figure. S5.**
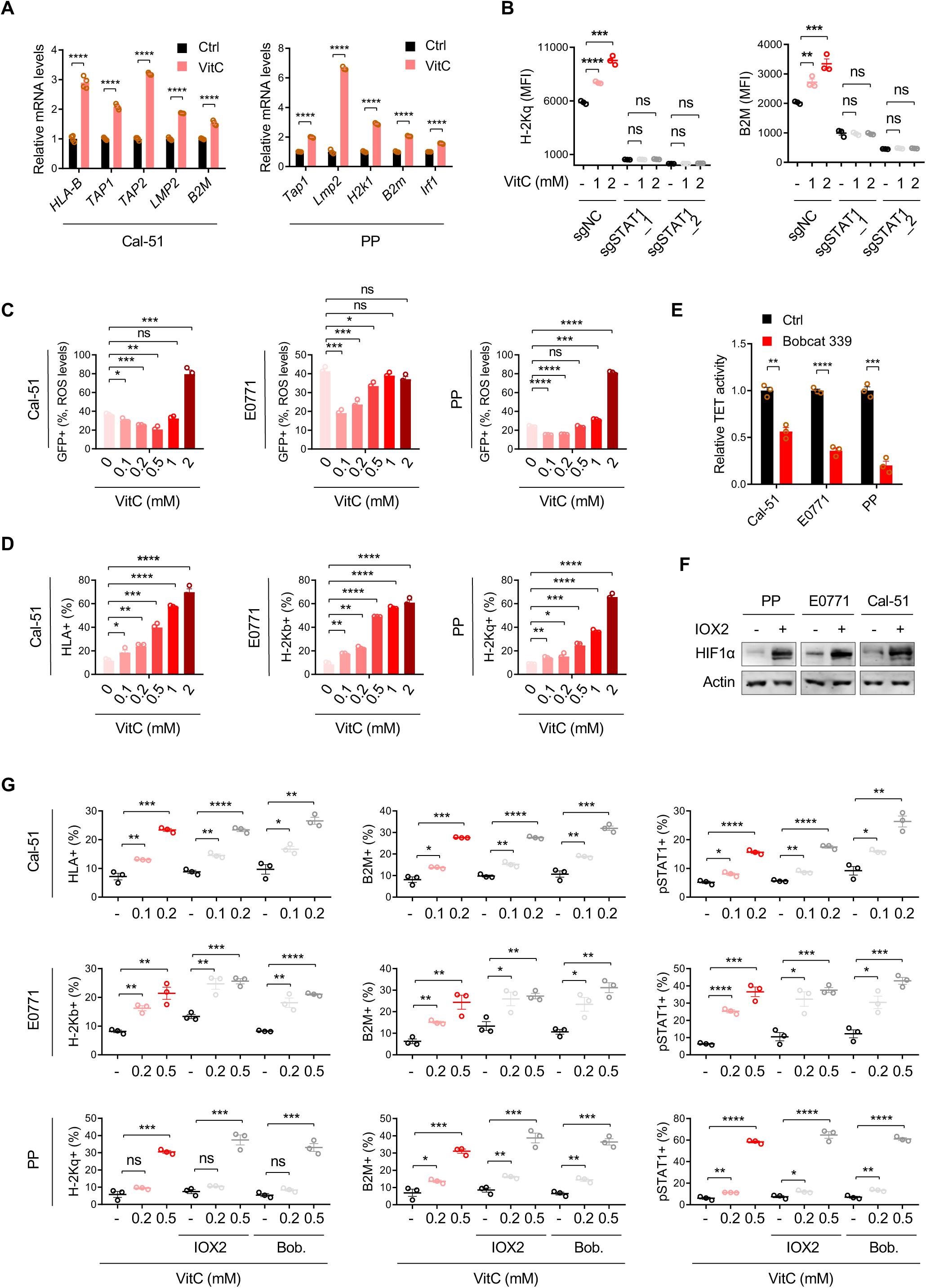
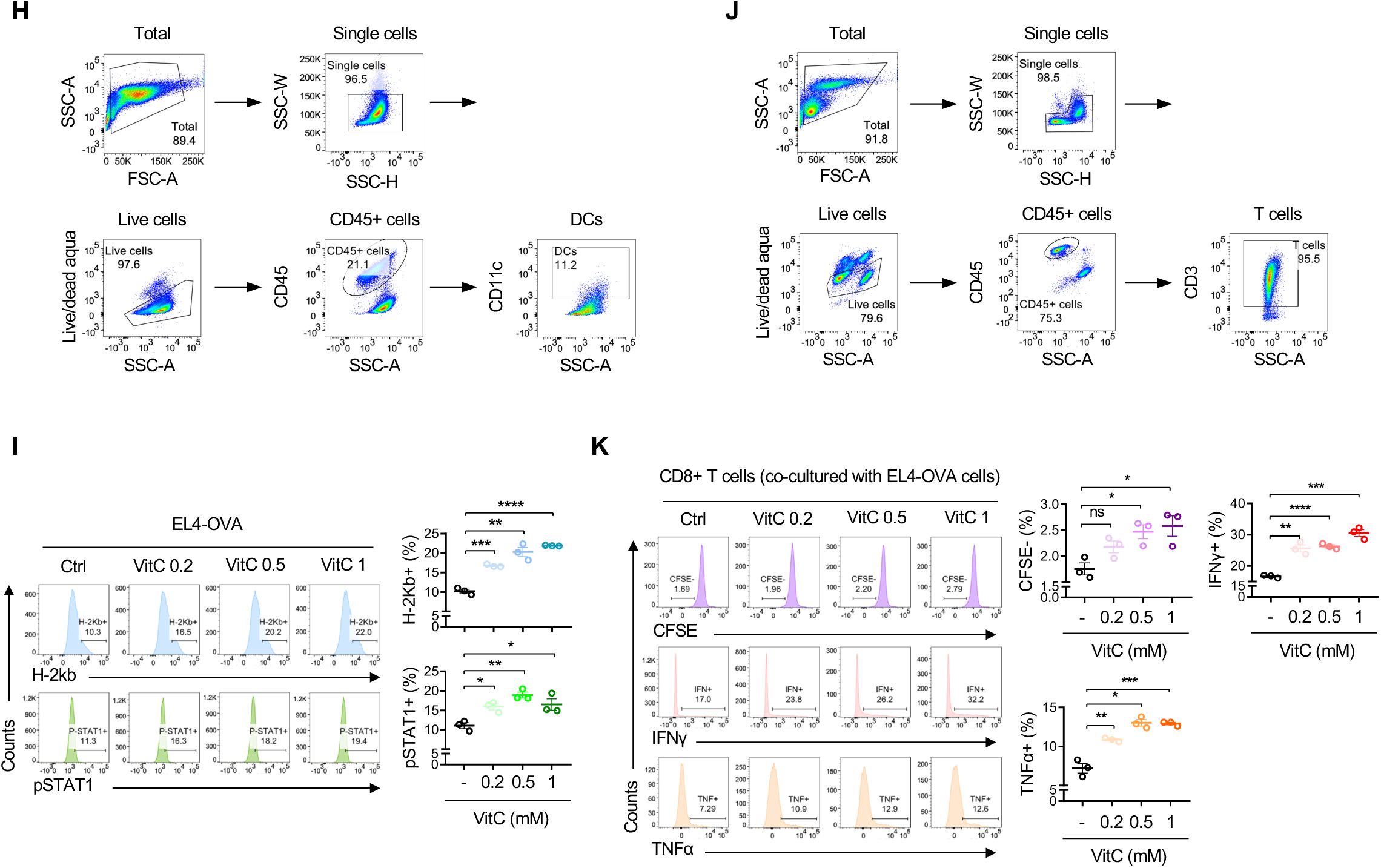
STAT1 K298 vitcylation facilitates MHC/HLA class I expression, related to Figure 5. (A) Quantitative PCR analysis of antigen processing and presentation genes expression in Cal-51 cells (n = 4) and PP cells (n = 3) treated with either vehicle or vitC for 2 days (0.5 mM vitC for Cal-51 cell treatment, 2 mM vitC for PP cell treatment). Data are represented as mean ± SEM. ****p < 0.0001. (B) Flow cytometry analysis of MHC class I expression in PP-sgControl, PP-sgSTAT1_1, and PP- sgSTAT1_2 cells cultured in different concentrations of vitC for 2 days (n = 3). Data are represented as mean ± SEM. **p < 0.01, ***p < 0.001, ****p < 0.0001. (C) ROS levels were measured in Cal-51, E0771, and PP cells treated with various concentrations of vitC for 2 days (n = 3). Data are represented as mean ± SEM. *p < 0.05, **p < 0.01, ***p < 0.001. (D) Flow cytometric analysis of MHC/HLA class I expression on Cal-51, E0771, and PP cells treated with different concentrations of vitC for 2 days (n = 3). Data are represented as mean ± SEM. *p < 0.05, **p < 0.01, ***p < 0.001, ****p < 0.0001. (E) TET activity was measured in Cal-51, E0771, and PP cells treated with vehicle or 10 μM Bobcat 339 for 6 hours (n = 3). Data are represented as mean ± SEM. **p < 0.01, ***p < 0.001, ****p < 0.0001. (F) HIF1α level was measured in Cal-51, E0771, and PP cells treated with vehicle or 0.2 μM IOX2 for 6 hours. (G) Flow cytometric analysis of pSTAT1 and MHC/HLA class I expression in Cal-51, E0771, and PP cells cultured in medium supplemented with the vehicle, vitC (48 hours), 10 μM Bobcat 339 (52 hours), vitC + 10 μM Bobcat 339, 0.2 μM IOX2 (52 hours) or vitC + 0.2 μM IOX2 (n = 3). Data are represented as mean ± SEM. *p < 0.05, **p < 0.01, ***p < 0.001, ****p < 0.0001. (H) Flow cytometry gating strategy for the DCs population. Hereafter for DCs gating. Representative plots are shown. (I) Flow cytometric analysis of H-2Kb and pSTAT1 expression on EL4-OVA cells treated with different doses of vitC for 3 days (n = 3). Data are represented as mean ± SEM. *p < 0.05, **p < 0.01, ***p < 0.001, ****p < 0.0001. (J) Flow cytometry gating strategy for T cells population. Hereafter for T cells gating. Representative plots are shown. (K) Flow cytometric analysis of CD8^+^ T (OT-I) cells co-cultured with EL4-OVA cells pretreated with 2 mM vitC. T cells (CD45^+^ CD3^+^ CD8^+^) proliferation and activity were quantified as CFSE^-^ and IFNγ^+^, TNFα^+^ cells, respectively (n = 3). Data are represented as mean ± SEM. *p < 0.05, **p < 0.01, ***p < 0.001, ****p < 0.0001.

**Figure. S6.**
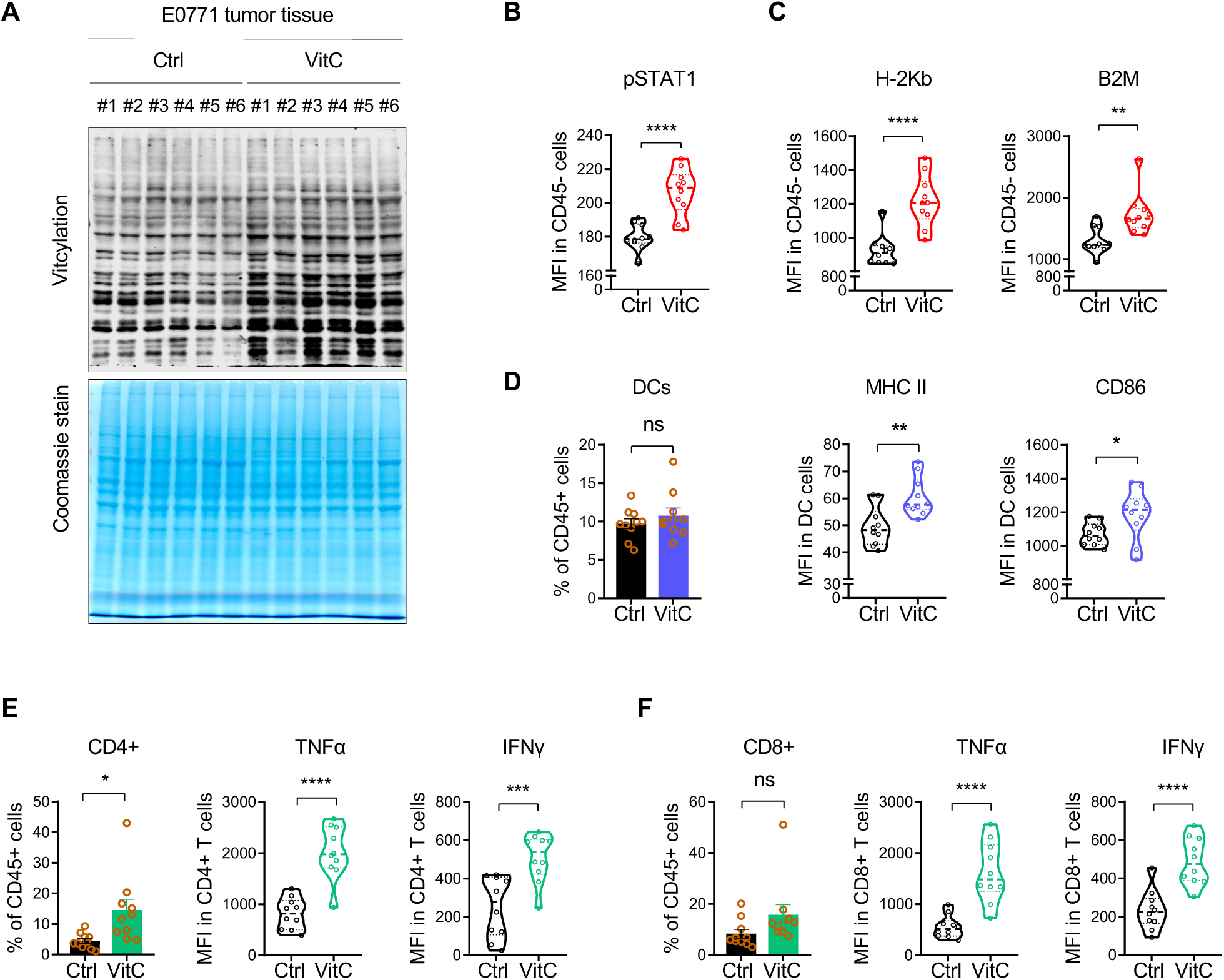
VitC treatment modulates the immune milieu *in vivo,* related to Figure 6. (A) Vitcylation levels in E0771 tumors were measured by WB with anti-vitcylation antibody (n = 6 for each group). Protein levels were normalized by coomassie staining. (B and C) Flow cytometry analysis of pSTAT1 (B), H-2Kb and B2M (C) expression on E0771 tumor cells (CD45^-^) from vehicle- or vitC-treated mice (vehicle n = 10, vitC treated n = 10). Data are represented as mean ± SEM. **p < 0.01, ****p < 0.0001. (D) Flow cytometry analysis of DCs (CD45^+^ CD11c^+^) population, and the MHC II and CD86 expression in DCs in E0771 tumor tissue from vehicle- or vitC-treated mice (vehicle n = 10, vitC treated n = 10). Data are represented as mean ± SEM. *p < 0.05, **p < 0.01. (E) Flow cytometry analysis of CD4^+^, CD4^+^ TNFα^+^, and CD4^+^ IFNγ^+^ T cells (CD45^+^ CD3^+^) isolated from E0771 tumors tissue from vehicle- or vitC-treated mice (vehicle n = 10, vitC treated n = 10). Data are represented as mean ± SEM. *p < 0.05, ***p < 0.001, ****p < 0.0001. (F) Flow cytometry analysis of CD8^+^, CD8^+^ TNFα^+^, and CD8^+^ IFNγ^+^ T cells (CD45^+^ CD3^+^) isolated from E0771 tumors tissue from vehicle- or vitC-treated mice (vehicle n = 10, vitC treated n = 10). Data are represented as mean ± SEM. ****p < 0.0001.

